# HBV DNA is a substrate for the cGAS/STING pathway but is not sensed in infected hepatocytes

**DOI:** 10.1101/867440

**Authors:** Lise Lauterbach-Rivière, Maïwenn Bergez, Saskia Mönch, Bingqian Qu, Maximilian Riess, Florian W. R. Vondran, Juliane Liese, Veit Hornung, Stephan Urban, Renate König

## Abstract

HBV chronic infection is a critical risk factor for hepatocellular carcinoma. Although debated, the absence of innate immune response to HBV infection in hepatocytes is becoming the current view. However the underlying reasons are poorly understood. This study aims to define potential viral pathogen-associated molecular patterns (PAMPs) and the pattern recognition receptors (PRRs), and to elucidate whether HBV counteracts the innate pathways.

The innate immune response to HBV infection was monitored by interferon-stimulated gene 54 (ISG54) mRNA, a direct downstream transcriptional target of Interferon Regulatory Factor 3 (IRF3), or IRF3 phosphorylation. The immunostimulatory potential of naked HBV DNAs or RNAs and the respective PRRs were determined upon viral nucleic acid transfection in immunocompetent cells including knockout cells lacking key molecules of innate pathways. The expression and functionality of DNA and RNA sensing pathways in primary human hepatocytes (PHH) were assessed. The inhibition of the DNA-sensing pathway by HBV was tested using IRF3 nuclear translocation assay.

Our study revealed that HBV infection does not induce an innate response in infected hepatocytes, even in absence of HBV X protein. HBV relaxed-circular DNA (rcDNA) and DNA replication intermediates, but not HBV RNAs, are immunostimulatory and sensed by Cyclic Guanosine Monophosphate-Adenosine Monophosphate Synthase (cGAS) and Stimulator of Interferon Genes (STING). Although PHH express DNA sensors to reduced levels compared to myeloid cells, they can respond to naked HBV rcDNA. However, we show that the absence of innate response to HBV infection in hepatocytes is not due to an active inhibition of the DNA sensing pathway by the virus.

HBV passively evades the innate immune response in infected hepatocytes by (i) producing non-immunostimulatory RNAs, (ii) avoiding sensing of its DNAs by cGAS/STING without active inhibition of the pathway, possibly through shielding of the viral DNAs by the capsid.

**Author summary:** Innate immune responses are the first line of defense against viral infections. They lead to the production of antiviral factors after recognition of specific viral features by the infected cells. Here we show that HBV, a major cause of liver cirrhosis and cancer, avoids recognition by infected hepatocytes through different means. First, HBV RNAs, contrarily to other viral RNAs, are not immunostimulatory. Second, we show that naked HBV DNAs are recognized by cGAS/STING and induce an innate immune response. Furthermore, we demonstrate that this pathway is active in hepatocytes and is not inhibited by the virus. Instead, we propose that HBV DNAs are not accessible to cGAS/STING in the context of an infection. This might be due to shielding of the viral DNA by the viral capsid.

## Introduction

Innate immunity is the first line of defense against pathogens. Detection of pathogen-associated molecular patterns (PAMPs) by cellular pathogen recognition receptors (PRRs) triggers signaling pathways leading to interferon (IFN) production, which induces antiviral interferon-stimulated genes (ISGs) and pro-inflammatory cytokines.

In viral infections, viral RNA or DNA are common PAMPs that can be detected by cytosolic PRRs. Members of the RIG-I-like Receptor (RLR) family such as Retinoic Acid Inducible Gene I (RIG-I) and Melanoma Differentiation-Associated protein 5 (MDA5) sense foreign RNAs and activate Mitochondrial Antiviral Signaling Protein (MAVS), while foreign viral DNA is recognized by sensors like cyclic GMP-AMP Synthase (cGAS). Activated cGAS produces 2’3’-cyclic GMP-AMP (cGAMP), which activates Stimulator of Interferon Genes (STING). STING or MAVS activation can both lead to the activation of Interferon Regulatory Factor 3 (IRF3), which promotes IFN genes transcription [1]. To evade antiviral innate responses, viruses have evolved escape strategies involving the inhibition of innate signaling pathways by viral proteins, or the shielding of the viral genome from innate sensors (reviewed in [1, 2]).

To date, more than 250 million people are chronically infected with HBV which is a high-risk factor for liver cirrhosis and hepatocellular carcinoma. Its interplay with the signaling pathways leading to IFN production is still a matter of debate. Some publications described a type I and III IFN or pro-inflammatory response to HBV infection in cultured hepatocytes [3–6]. HBV may also induce pro-inflammatory cytokines in immune cells such as Kupffer cells, the liver macrophages, even if they are not productively infected [7–9]. The induction of an innate response to HBV in infected hepatocytes has however been questioned by several recent reports [8,10–13], confirming initial studies in chimpanzees [14] and patients [15, 16]. Therefore, HBV was proposed to be a stealth virus in infected hepatocytes, and the mechanisms behind the lack of innate responses are not fully understood.

HBV particles contain a partially double-stranded DNA, the relaxed-circular DNA (rcDNA), which is repaired into a covalently closed circular DNA (cccDNA) in the nuclei of infected cells. The cccDNA is then transcribed into mRNAs and pregenomic RNA (pgRNA) that are exported to the cytoplasm. The pgRNA is encapsidated into newly synthesized capsids, where it is reverse transcribed into rcDNA. Both HBV RNAs and DNAs are therefore potential PAMPs. HBV pgRNA has been proposed to be sensed by the RIG-I- [4] or MDA5-pathways [3]. Surprisingly, MDA5 was also reported to bind HBV DNA [3]. Recently, sensing of HBV DNA by the cGAS/STING pathway was suggested [17, 10] but the expression and functionality of this pathway in hepatocytes is unclear [8,10,13,17,18]. Therefore, the immunostimulatory potential of HBV nucleic acids and the PRR involved require further investigations.

HBV could inhibit the innate immune response to escape its antiviral effects, as suggested by several publications [6,10,19–28]. In particular, the regulatory Hepatitis B virus X protein (HBx) has been described to inhibit the MAVS pathway [20, 26], while the viral polymerase was reported to inhibit STING [29] and the phosphorylation of IRF3 [23, 24]. Verrier et al. [10] suggested the down-regulation of cGAS, STING and TANK binding kinase 1 (TBK1) by HBV but did not show whether this down-regulation had an effect on the functionality of the pathway [10]. On the other hand, several publications claimed an absence of inhibition of the innate response by HBV. Most of them tested several RNA-sensing pathways [8,30,31], but the DNA-sensing pathway has been less extensively investigated. Guo et al. [11] proposed that HBV expression does not affect the response to dsDNA or cGAMP stimulation in HepAD38 cells expressing an integrated copy of HBV genome under the control of a tetracycline-controlled promoter, but did not study a genuine HBV infection.

Here, we confirmed the lack of innate response to HBV infection in hepatocytes, which was not restored by HBx depletion. To understand the lack of innate response to HBV infection we first assessed the immunostimulatory potential of naked HBV DNAs and RNAs in monocyte-derived dendritic cells (MDDCs) as a model for highly immunocompetent cells. For the first time, we could demonstrate that HBV RNAs are not immunostimulatory. On the contrary, naked HBV rcDNA can elicit a strong innate response, mediated by the cGAS/STING pathway. In hepatocytes, this pathway is expressed at a low level but retains its ability to sense HBV rcDNA. Furthermore, we demonstrate that HBV infection does not inhibit the innate immune response to foreign DNA in hepatocytes.

## Results

### HBV does not induce an innate immune response in hepatocytes

We first determined whether hepatocytes could mount an innate immune response to HBV infection. HepG2-hNTCP cells, overexpressing the viral receptor NTCP, were infected with HBV for 16 days. ISG54 mRNA was chosen as a marker for the innate immune response, as it is a direct target gene of IRF3-dependent transcription [32, 33]. IFN-λ1 mRNA expression, previously proposed to be induced by HBV [4, 5] was also analyzed (Fig 1). Although the cells were efficiently infected, as demonstrated by the accumulation of HBV RNAs over time, we observed no induction of ISG54 or IFN-λ1 (Fig 1A). The innate response to HBV infection was next investigated in primary human hepatocytes (PHH) (Fig 1B). HBV RNAs could be detected 4 dpi, demonstrating the efficiency of infection. However, no ISG54 was induced in HBV-infected PHH. As a positive control, infection with Sendai virus (SeV), an RNA virus, induced an innate response in HepG2-hNTCP and PHH, indicating that they are competent for cytosolic sensing.

**Fig 1:**
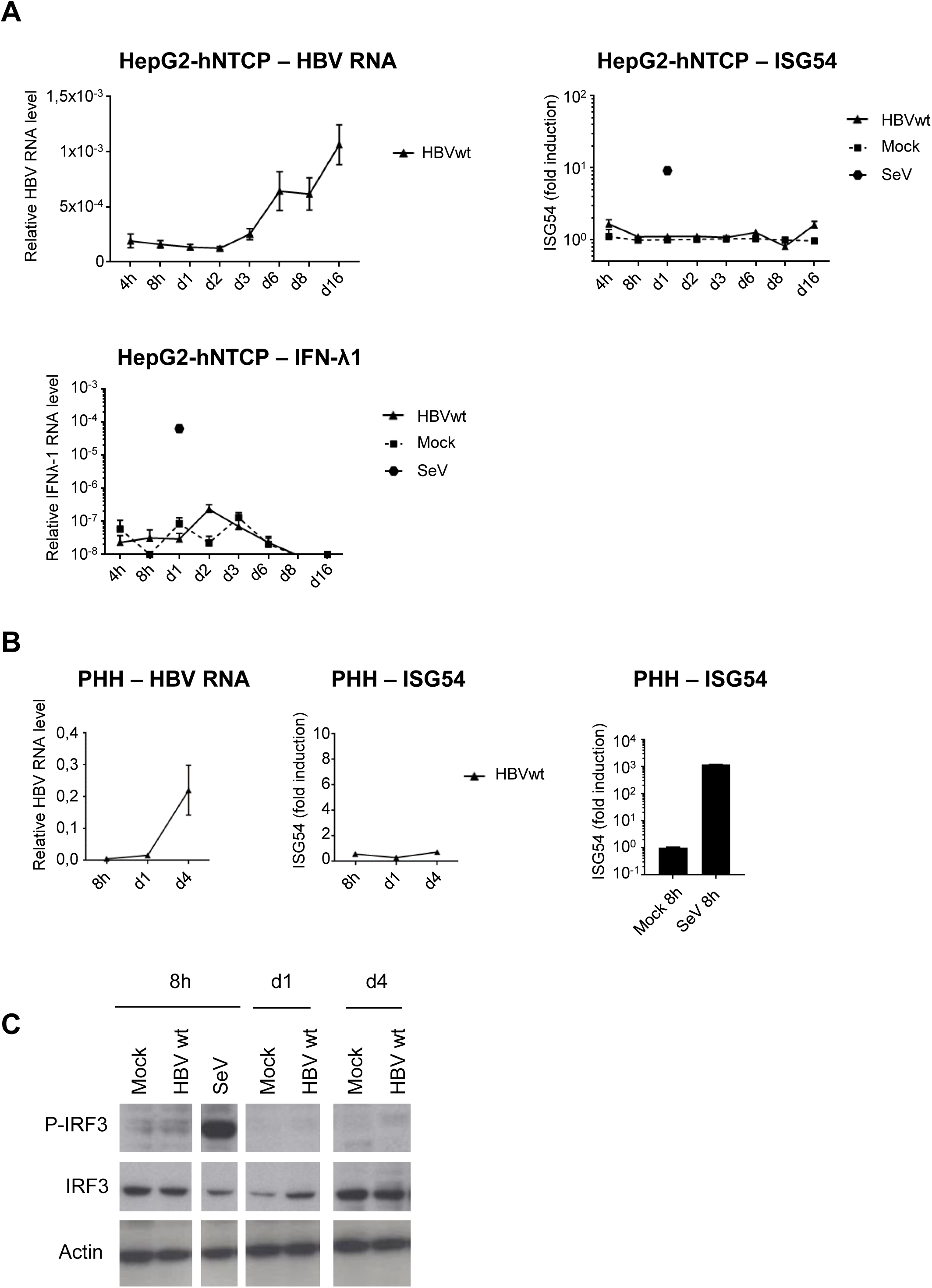
HBV does not induce an innate response in hepatocytes. HepG2-hNTCP (A) or PHH (B, C) were infected with HBVwt (MOI 100) or SeV (MOI 0,13). HBV, ISG54 or IFN-λ1 RNAs were quantified by RT-qPCR (A, B). Relative expression levels to the reference gene RPL13A were calculated. For ISG54, fold induction to the respective mock-control is shown for each time point. Average and standard error of the mean (SEM) of 3 independent (HepG2-hNTCP) or 1 (PHH) experiments in technical triplicates are shown. (C) Phospho- or total IRF3 protein levels were analyzed by Western blot in HBVwt-infected PHH (MOI 100).

Since HBx may inhibit innate sensing pathways [20, 26], we used an HBx-defective mutant of HBV (HBV X-). HBV X- has been previously shown to enter hepatocytes and efficiently form cccDNA. HBV X-RNA transcription rate is reduced compared to HBVwt, but not totally abolished [34], allowing to use this mutant to study the innate response in infected hepatocytes. Even in the absence of HBx, no innate response was observed in infected HepG2-hNTCP (S1A Fig) or PHH (S1B Fig).

We next investigated IRF3 activation by phosphorylation, as an upstream event common to both RNA and DNA cytosolic sensing pathways. No IRF3 phosphorylation was detected upon HBVwt (Fig 1C) or HBV X- (S1C Fig) infection of PHH, while SeV infection induces a strong IRF3 phosphorylation after 8h (Fig 1C, and S7 Fig).

Altogether, our data confirm the absence of an innate response to HBV infection in hepatocytes and rule-out an inhibition of this response by HBx. Furthermore, the absence of IRF3 phosphorylation suggests either an absence of sensing by PRRs or, alternatively, an inhibition of the signaling pathways upstream of IRF3 phosphorylation.

### HBV DNA but not RNA is immunostimulatory

To understand the lack of innate response to HBV infection, we next determined whether HBV nucleic acids (DNAs or RNAs) are immunostimulatory. To this aim, we used monocyte-derived dendritic cells (MDDCs) as a model for highly sensitive immune cells that most likely express all nucleic acid sensors. The MDDCs were transfected with 10-fold serial dilutions of quantified viral nucleic acids and ISG54 mRNA induction was measured by RT-qPCR. The copy number of the viral nucleic acids (DNA or RNA) used for MDDC transfection was determined as described in the material and methods section and is indicated in Table 1.

**Table 1.** Quantification of viral nucleic acids used for MDDC transfection experiments.

|  | SeV n.ac.<br>from<br>particles | MVA n.ac.<br>from<br>particles | HBV n.ac.<br>from<br>particles | HepaD38<br>RNA | HBV repl.<br>intermediate<br>s |
| --- | --- | --- | --- | --- | --- |
| **Viral DNA (copies/cell)** |  | 5.0E+03 | 5.0E+03 |  | 5.0E+03 |
| **Viral RNA (cDNA-**<br>**equivalent copies/cell)** | 2.8E+01 |  | 2.3E+01 | 1.3E+04 |  |

(n.ac: nucleic acids; repl. intermediates: replication intermediates)

To validate the system, we first used total nucleic acids isolated from viral preparations of the RNA virus SeV or of a Gfp-expressing vector based on the DNA virus Modified vaccinia Ankara (MVA-gfp). SeV RNA activates RIG-I signaling and is used as a model to stimulate cytoplasmic sensing of RNA [35]. MVA-gfp DNA activates cGAS/STING signaling and is used as a model to stimulate cytoplasmic sensing of DNA [36]. We observed a high ISG54 induction upon transfection of MDDCs with SeV nucleic acids (Fig 2) demonstrating the functionality of the RIG-I pathway. This response is abrogated upon RNase digestion of SeV nucleic acids, while it is unaffected by DNAse digestion, proving that this response is specific for RNA. In this assay, the RIG-I pathway is able to respond to as little as 2,8E-1 cDNA-equivalent copies of SeV RNA per cell (Fig 2, 1:100 dilution of SeV nucleic acids quantified in Table 1). Nucleic acids from MVA-gfp particles also induce a dose-dependent ISG54 response, validating the functionality of the cGAS/STING pathway with a detection limit below 5E+2 copies of MVA-gfp DNA/cell (Fig 2, 1:10 dilution of MVA-gfp nucleic acids quantified in Table 1).

**Fig 2:**
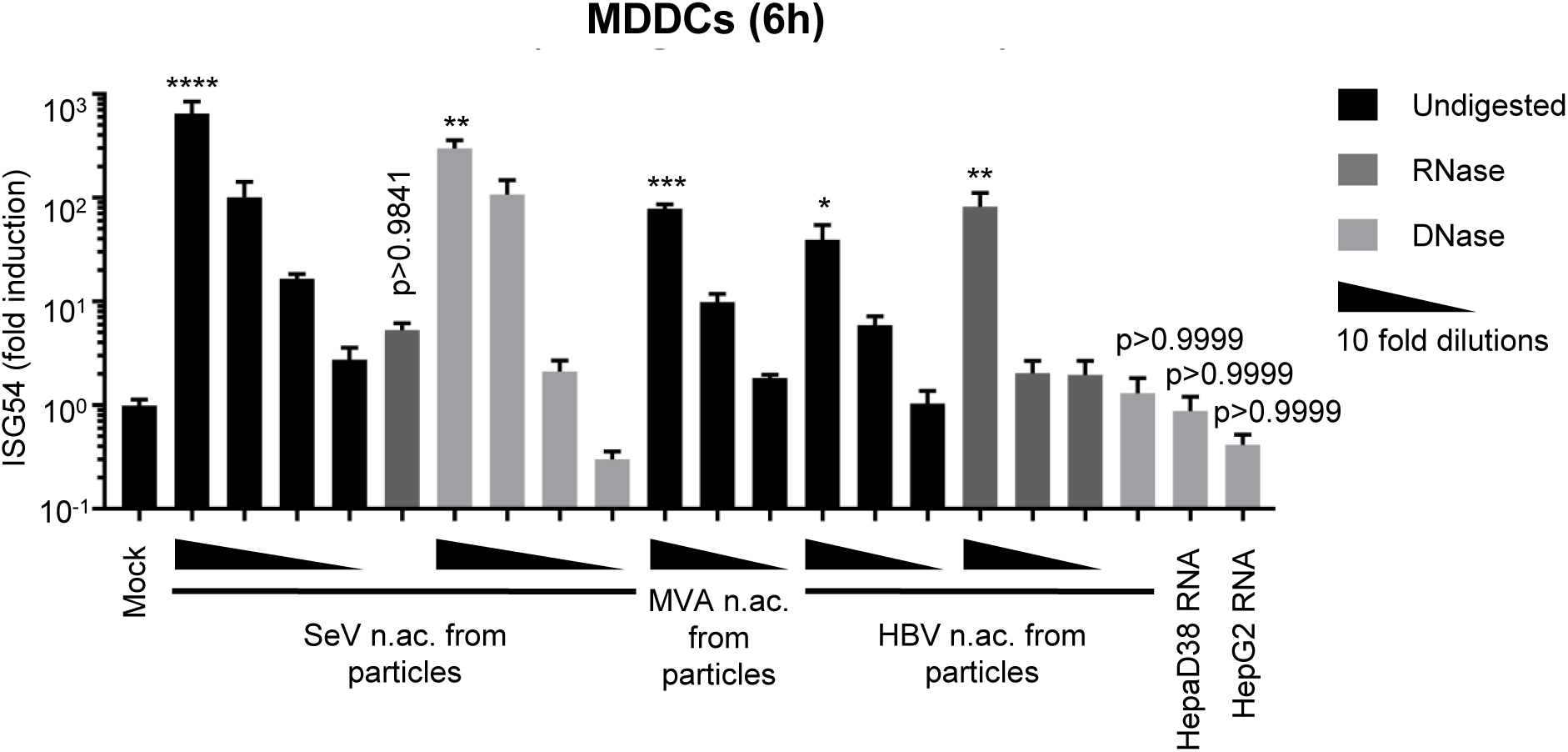
HBV DNA but not RNA is immunostimulatory. Total nucleic acids (n.ac.) were extracted from SeV, MVA-gfp or HBV particles. Total RNAs were extracted from HepAD38 or HepG2 cells. When indicated, the nucleic acids were digested with RNase (dark grey) or DNase (light grey; black bars: undigested). Human MDDCs were transfected with 10-fold dilutions of the nucleic acids (quantified in Table 1). ISG54 mRNA level was quantified by RT-qPCR 6h post-transfection. Average and SEM of 3 technical replicates of at least 2 donors are shown. Levels of significance: ****p<0,0001, ***p<0,001, **p<0,01 and *p<0,05 (Kruskal-Wallis-Test with Dunn’s multiple comparisons test).

Interestingly, nucleic acids extracted from HBV viral preparations induce an ISG54 response, which is unaffected by RNase digestion but is abrogated by DNAse digestion (Fig 2), showing that HBV relaxed circular DNA (rcDNA) contained in the viral particles is immunostimulatory. HBV viral preparations also contained small amounts of HBV RNAs (Table 1), confirming the reports by Cheng et al. [8]. However, these RNAs cannot account for the innate response in transfected MDDCs since the response is abrogated by DNase digestion (Fig 2).

Furthermore, we tested the immunostimulatory potential of HBV RNAs using total RNAs from HepAD38 cells. HepAD38 contain an integrated copy of HBV genome and produce all viral RNAs [37]. However, no ISG54 induction was detected upon transfection of as many as 1,3E+4 cDNA-equivalent copies/cells of HBV RNAs (Fig 2, Table 1), which is 4,6E+4 times above the detection limit for SeV RNAs in this MDDC stimulation assay (2,8E-1 copies/cell). This indicates that HBV RNAs (mRNAs or pgRNA) are not immunostimulatory. In HBV-infected cells, various DNA replication intermediates are produced in the cytoplasm upon reverse-transcription of the pgRNA [38] and may also act as PAMPs. To test their immunostimulatory potential, we extracted HBV DNA replication intermediates from the cytoplasmic fraction of HepAD38 cells. Intriguingly, they induced an ISG54 response in transfected MDDCs, however, the response was slightly weaker than when using the same copy number (5000 copies/cell) of HBV rcDNA (S2 Fig). In summary, naked HBV rcDNA and DNA replication intermediates elicit an innate response whereas HBV RNA species are not immunstimulatory.

### HBV DNA is sensed by the cGAS/STING pathway

To identify which PRRs and pathways sense and respond to HBV rcDNA, we used a panel of THP-1 knock-out (KO) cell lines deficient for key nodes of the sensing pathways, cGAS, STING or MAVS (THP-1 wt, ΔSTING, ΔcGAS and ΔMAVS respectively, Fig 3A and S8 Fig). As expected, KO of STING or cGAS did not significantly affect the ISG54 response to SeV infection, while MAVS KO abrogated it (Fig 3B). On the contrary, STING or cGAS KOs abrogated the innate response to transfection with the cGAS agonist herring testes DNA (HT-DNA) while the transfected DNA was well sensed by MAVS KO, proving the validity of the chosen assay system. Similar to MDDCs, transfection of HBV nucleic acids from viral particles in THP1 wt strongly induced a dose-dependent ISG54 response, which is abrogated by DNAse digestion. Interestingly, STING and cGAS KOs totally abrogated the response to HBV nucleic acids, while MAVS KO had no significant effect. These results indicate that HBV rcDNA is sensed through the cGAS/STING pathway, while the RLR pathway is not involved.

**Fig 3:**
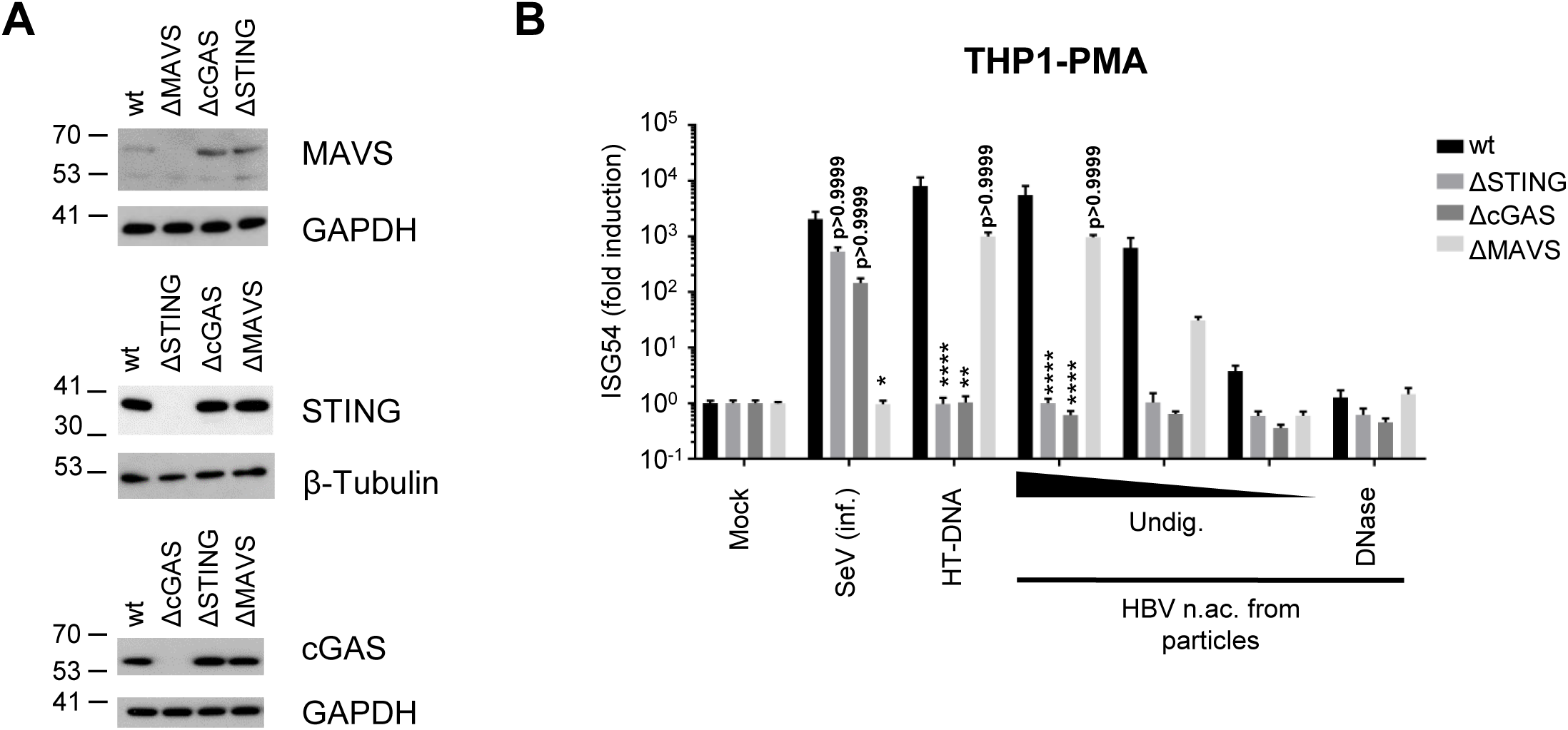
HBV rcDNA is sensed by the cGAS/STING pathway. (A) Genome editing of THP1 CRISPR/Cas9 KO for STING, cGAS or MAVS was controlled by Western Blot. (B) PMA-differentiated WT or KO THP1 cells were infected with SeV, transfected with HT-DNA (0,1 µg/well) or with undigested (Undig.) or DNAse-digested HBV nucleic acids (n.ac.) from particles. ISG54 mRNA fold induction to the mock was determined by RT-qPCR 24h post-transfection. Average and SEM of 3 independent experiments in triplicates are shown. ****p<0,0001, **p<0,01 and *p<0,05 (Kruskal-Wallis-Test with Dunn’s test; For each stimulus, the knock-out cells are compared to the WT cells).

### Hepatocytes express low level of the DNA sensors compared to immune cells

Having proven that HBV rcDNA is able to stimulate the cGAS/STING pathway, we wondered whether the absence of innate response to HBV infection in hepatocytes is due to a deficiency of this pathway in hepatocytes. To this aim, we first measured by RT-qPCR the RNA level of cGAS, STING as well as other components of PRR pathways in hepatic cell lines and in PHH and compared them to primary immune cells (MDDCs, monocyte-derived macrophages (MDM), Kupffer cells) and THP1 cells (Fig 4). cGAS RNAs are expressed to a similar level in all the tested immune cells, however its expression is strongly reduced in hepatic cells although still detectable in PHH. STING shows a variable expression pattern in immune cells (high in THP1, MDMs and Kupffer cells, lower in MDDCs as seen earlier in [39]). In hepatic cells, STING expression is generally lower than in MDDCs, although always above the detection limit. Gamma-interferon-inducible protein 16 (IFI16), proposed as immune sensor of retroviral DNA intermediates and a cGAS cooperative factor for activating STING [40], is expressed in PHH and HepaRG-hNTCP, although slightly reduced compared to immune cells, but is undetectable in the HepG2 derived cell lines. PQBP1, a cofactor of the cGAS/STING pathway involved in sensing of HIV-1 reverse-transcribed DNA [41], is expressed to similar levels in all tested cell types. Finally, RIG-I and MAVS are expressed to similar levels in immune and hepatic cells. In conclusion, some components of the DNA sensing pathways, and in particular the cGAS/STING pathway are considerably less expressed in hepatocytes than in immune cells, while components of the RNA sensing pathways are expressed to comparable levels.

**Fig 4:**
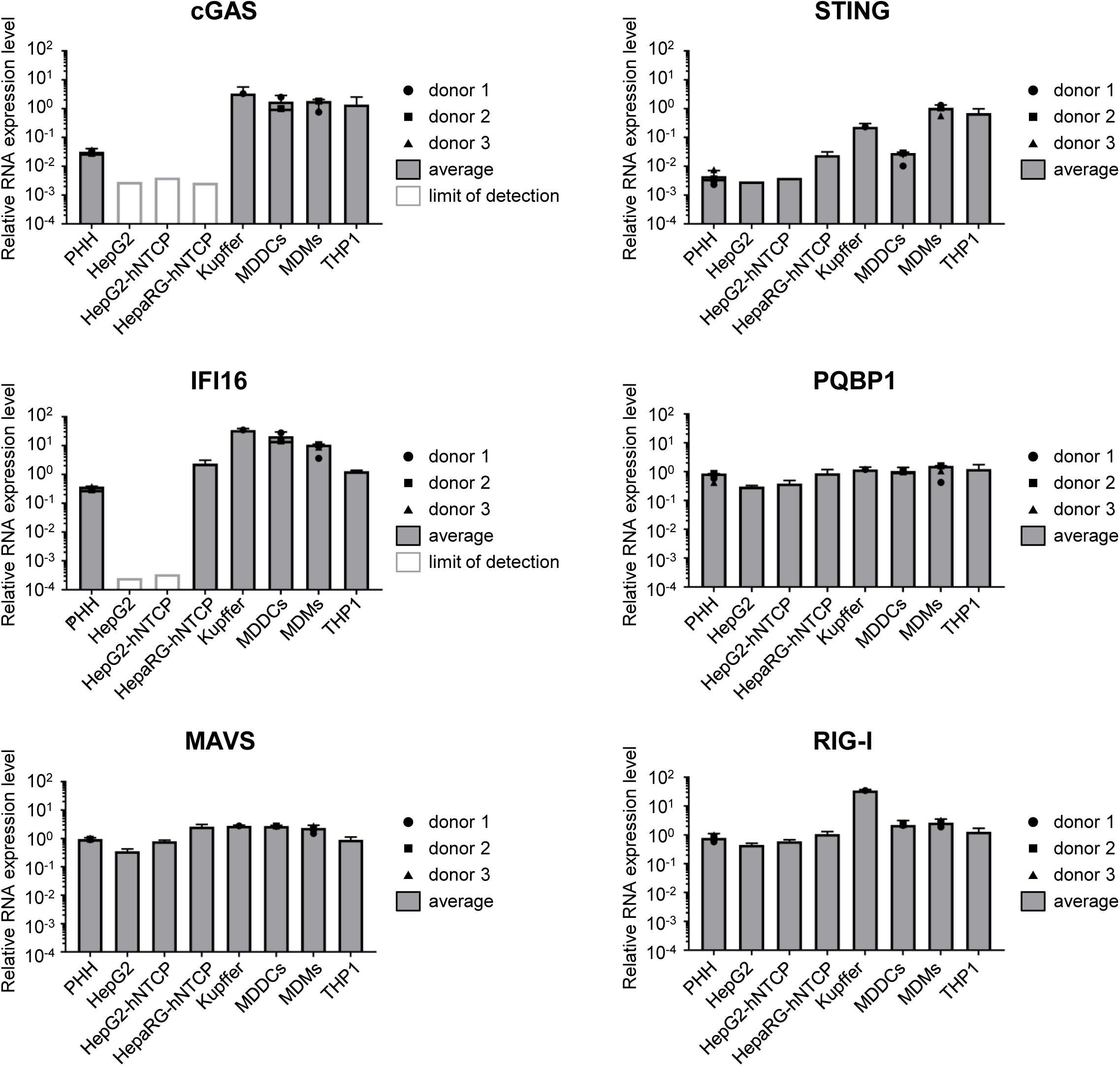
Hepatocytes express a low level of components of the DNA sensing pathway. cGAS, STING, IFI16, PQBP1, MAVS and RIG-I mRNAs were quantified by RT-qPCR in the indicated cell types and normalized to the geometric mean of RPL13A and TATBP mRNAs. For primary cells, the average and SEM of technical triplicates from 3 (PHH and MDMs), 2 (MDDCs) or 1 (Kupffer cells) donors are shown.

### Hepatocytes are competent for DNA sensing

We next tested whether the low levels of cGAS/STING correlate with an altered functionality of the pathway in hepatocytes. To this aim, we first stimulated HepG2-hNTCP with the STING agonist cGAMP (Fig 5A). Interestingly, ISG54 was transiently but significantly induced in HepG2-hNTCP 8 hours after cGAMP stimulation, indicating the functionality of STING and of the downstream pathway. We next tested the complete cGAS/STING pathway using infection with MVA-gfp, or transfection with HT-DNA and with HBV nucleic acids from viral particles. To determine if the levels of cGAS and STING are limiting for DNA sensing, we overexpressed cGAS and STING in HepG2-hNTCP using lentiviral vectors and compared with cells transduced with a control vector (Fig 5B). The infection efficiency of MVA-gfp was controlled by the GFP expression (S3 Fig). Since the transfection reagent alone induced a weak ISG54 response compared to an untransfected control, we compared the transfected samples to a mock-transfected control (transfection reagent only), while the infected samples were compared to an uninfected/untransfected control (mock). Interestingly, the control-transduced HepG2-hNTCP cells were able to respond to transfection with HT-DNA and HBV nucleic acids from viral particles, and to a lower extent, to MVA-gfp infection (Fig 5B). The response to HBV nucleic acids transfection was dose-dependent (1500 copies/cell induce ISG54 by 2 log, while 150 copies/cells are insufficient) and abrogated by DNAse digestion, demonstrating the specificity of the response for DNA. Transduction with cGAS and STING vectors increased the RNA level of cGAS and STING by 1 to 2 log compared to the control vector (S4 Fig) and enhanced the response to all DNA stimuli but not to SeV infection. This shows that HepG2-hNTCP are able to sense DNA, including HBV rcDNA, to some extent, but cGAS and STING expressions are limiting.

**Fig 5:**
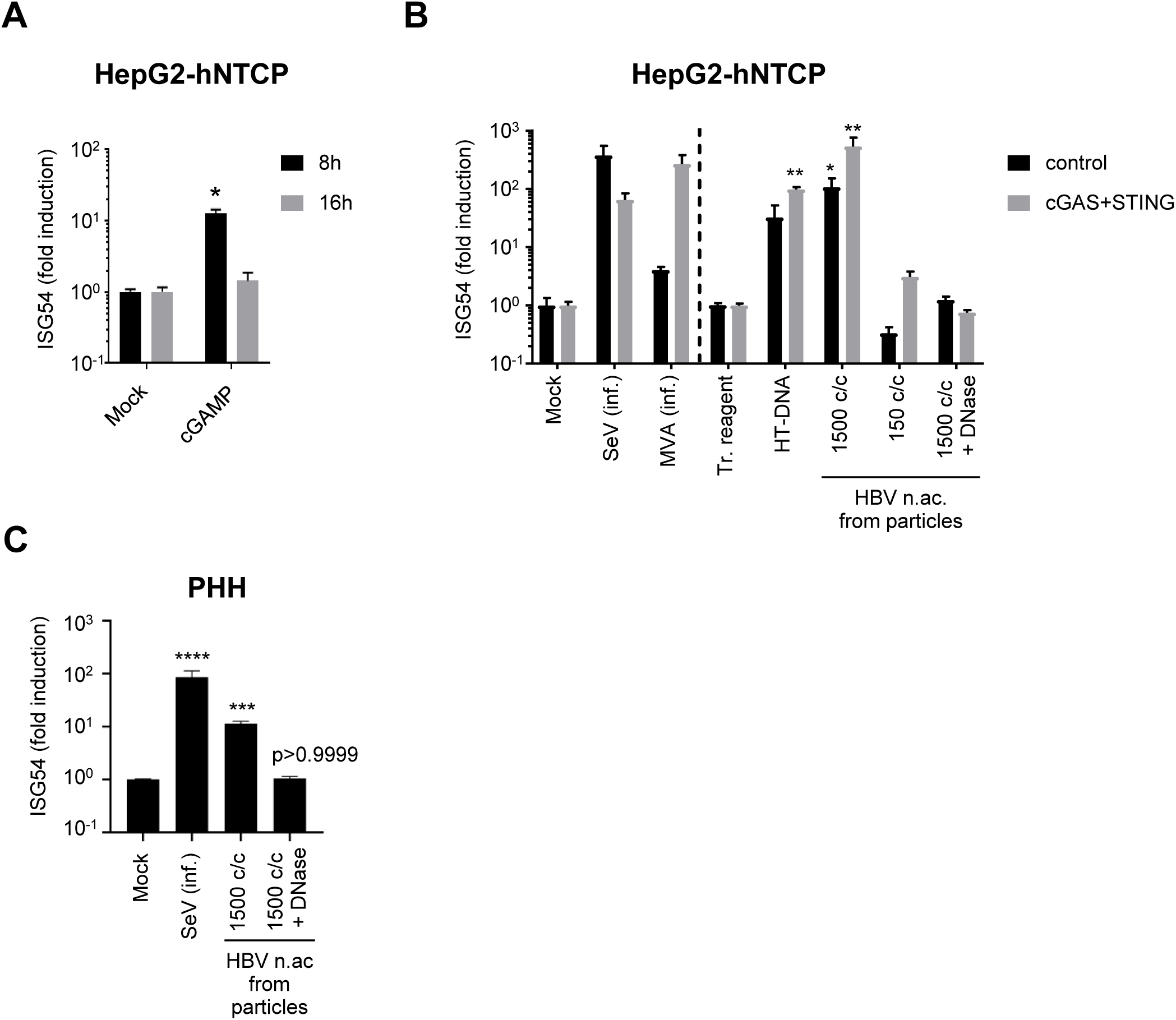
Hepatocytes respond to foreign DNA. HepG2-hNTCP (A), HepG2-hNTCP-control or -cGAS/STING (B) or PHH (C) were stimulated with cGAMP (A), infected (inf.) with SeV or MVA-gfp, or transfected with HT-DNA or HBV nucleic acids (c/c: copies/cells) (B, C). ISG54 mRNA fold inductions to mock (A, B left part, C) or to transfection reagent-treated cells (tr. reagent) (B, right part) was determined by RT-qPCR. Average and SEM of 4 (A), 2 (B) independent experiments in triplicates or of 4 donors in duplicates (C). ****p<0,0001, ***p<0,001, **p<0,01, *p<0,05 (2-way ANOVA with Sidak’s multiple comparisons (A); Kruskal-Wallis with Dunn’s test (B, C)).

We next assessed the functionality of the DNA sensing pathway in PHH (Fig 5C). Interestingly, transfection of PHH with undigested, but not with DNAse-digested HBV rcDNA (1500 copies/cell) induced ISG54 in all 4 donors, indicating that PHH are able to sense HBV rcDNA and that the response is specific for DNA.

### HBV does not inhibit the innate response to foreign DNA in hepatocytes

The absence of the innate response to HBV could be explained by the inhibition of the DNA sensing pathway by the virus. This hypothesis has been suggested by previous studies: an inhibition of STING and IRF3 phosphorylation by HBV polymerase has been reported [23, 29]. In addition, Verrier et al. [10] observed a down-regulation of cGAS and STING in HBV-infected cells but did not show whether this impairs the functionality of the pathway. On the contrary, Guo et al. [11] observed no inhibition of the innate response to dsDNA or cGAMP upon HBV expression in HepAD38 cells. We therefore addressed this controversial question in HBV-infected HepG2-hNTCP. In order to focus only on infected cells, we immunostained HBc as an infection marker, and analyzed IRF3 nuclear translocation after HT-DNA transfection. IRF3 is translocated to the nucleus upon stimulation of the cGAS/STING pathway. To get a higher percentage of IRF3 nuclear translocation, we used HepG2-hNTCP overexpressing cGAS and STING (S5 Fig). No nuclear staining was observed for IRF3 in the unstimulated cells, independent of HBV infection (Fig 6), consistent with the lack of innate response to HBV infection. In the mock samples, HT-DNA transfection induced IRF3 nuclear translocation in 16,4% of the cells (Fig 6B, graph). We observed a similar rate of IRF3 translocation in HBc-positive, HBV-infected cells transfected with HT-DNA (19,4%; the ratio +/- standard deviation of IRF3 nuclear translocation in HT-DNA-transfected HBV-infected cells/mock is 1,18 +/- 0,55), demonstrating that HBV infection does not inhibit the innate immune response to foreign DNA.

**Fig 6.**
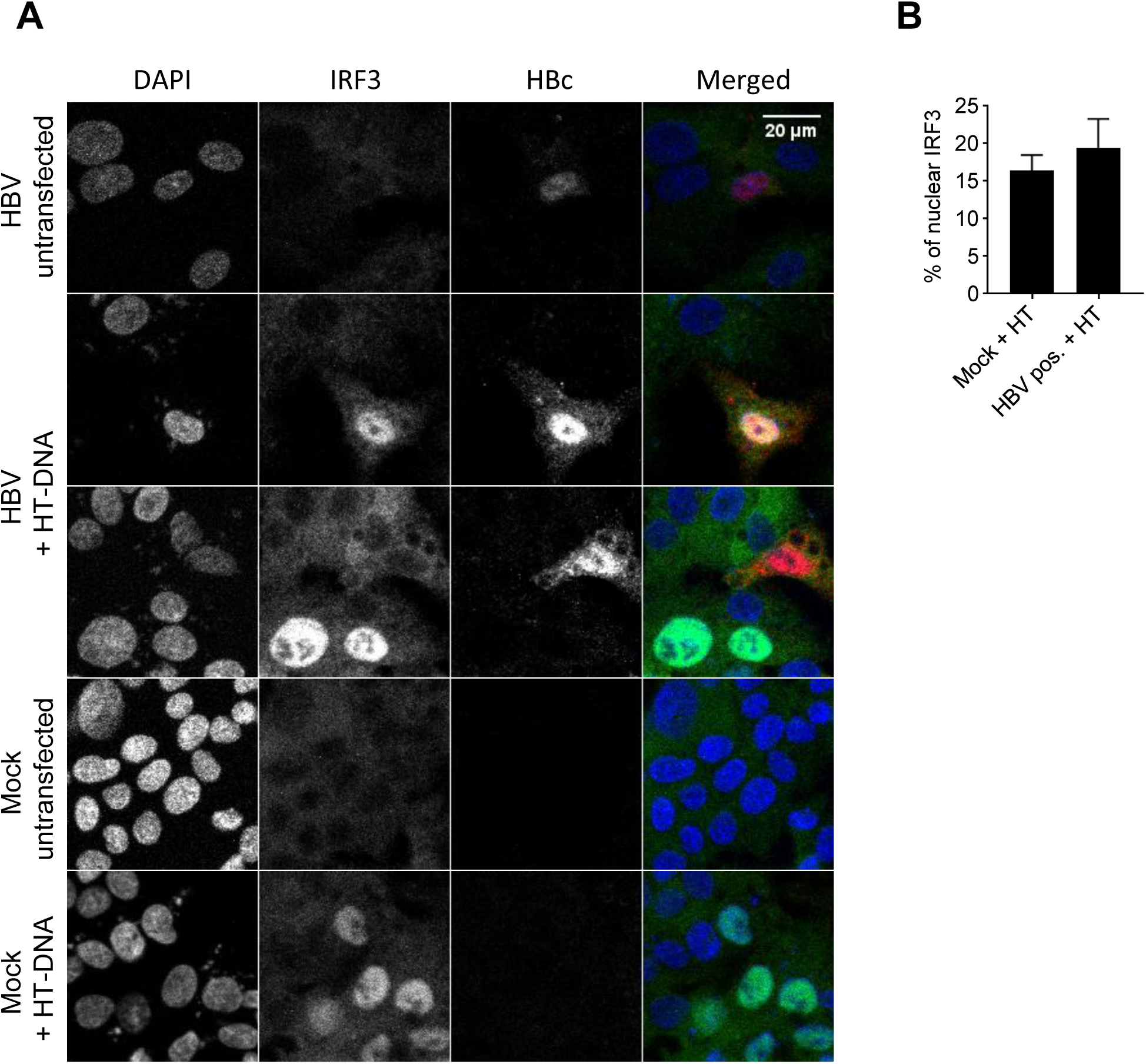
HBV does not inhibit the innate response to foreign DNA. HepG2-hNTCP-cGAS-STING were infected or not with HBVwt (MOI 10000). 7 dpi, the DNA-sensing pathway was stimulated with HT-DNA. 16 hours post-HT-DNA-transfection, IRF3 (green), HBc (red) and DNA (blue) were stained. (A) Representative images. (B) Percentage of nuclear IRF3 counted in HT-DNA transfected cells. For HBV-infected cells, the analysis is restricted to the HBc-positive cells (productive infection). Average and SEM of 2 independent experiments in duplicates are shown. At least 67 cells per condition and replicate were counted (total: 1153 mock-infected cells; 333 HBc-positive cells).

In addition, we measured the level of cGAS and STING mRNAs in mock-infected versus HBV-infected PHH (MOI 5000). Immunofluorescence staining of HBc indicated that the majority of the cells were infected (S6A Fig). However, productive HBV infection did not affect the levels of cGAS or STING compared to uninfected PHH or PHH infected in the presence of the entry inhibitor Myrcludex B (S6B and C Figs).

## Discussion

In this study, we confirmed the absence of an innate immune response to HBV infection in hepatocytes and investigated the reasons for this escape. We first demonstrated that naked HBV RNAs are not immunostimulatory. In contrast, we found that naked HBV DNAs, and particularly the rcDNA from viral particles, can be sensed by the cGAS/STING pathway. We showed that in hepatocytes, including PHH, this pathway is expressed at a low level, but is sufficient to respond to transfected HBV rcDNA. We further demonstrated that HBV does not actively inhibit the innate response to foreign DNA in infected hepatocytes.

We observed no induction of ISG54 or IFN-λ1 and no phosphorylation of IRF3 in HBV-infected hepatocytes (Fig 1). These data differ from Shlomai et al. [5] and Sato et al. [4], who detected a type I and/or III IFN response to HBV infection in hepatocytes, but are in line with the first observations made by Wieland et al. [14] in infected chimpanzees as well as with recent studies [8,10,12,31].

We showed for the first time that naked HBV RNAs, including mRNAs and the pgRNA, are not immunostimulatory in transfected MDDCs, used as a model for highly immune competent cells. These results are in contrast with Sato et al. [4], who reported that HBV pgRNA could be sensed by RIG-I, and Lu et al. [3], who proposed that HBV RNA could bind MDA5. Sato et al. suggested that the epsilon element (ε) of the pgRNA, that folds into a stem-loop structure, was the motif recognized by RIG-I. In their study, the genotype C of HBV was used, while we used the genotype D (ayw). However, the epsilon element of these two genotypes are identical, the difference in HBV genotypes is thus unlikely to explain the difference in our results. Cellular mRNAs bear specific features that avoid their recognition by the RLR sensors [42, 43]. Binding to RIG-I is synergistically inhibited by 7-methyl guanosine (Cap0) and 2’-O-methylation (Cap1) that cap human mRNAs [44]. All HBV RNAs are transcribed by the cellular RNA polymerase Pol II, similar to cellular mRNAs and have been described to bear a Cap0 [45, 46]. The presence of a Cap1 on HBV RNAs has not been formally studied, but could account for the absence of recognition by RIG-I. In addition, HBV RNAs contain N6-methyladenosine (m6A) RNA methylation [47], which has been shown to impair RIG-I binding [48].

On the contrary, we demonstrated that HBV rcDNA from viral particles is immunostimulatory in myeloid as well as hepatic cells including PHH. CRISPR-Cas9 knockout cells revealed that HBV rcDNA is sensed by the cGAS/STING pathway, while MAVS is dispensable, ruling out the possible sensing of HBV DNA by MDA5 suggested by Lu et al. [3]. Our results confirm and extend earlier studies showing the role of the cGAS/STING pathway in HBV DNA sensing [10, 17]. In addition, we demonstrated that HBV replication intermediates are immunostimulatory as well, although to a lesser extent than rcDNA.

The activity of the cGAS/STING pathway in hepatocytes is controversial. Several publications indicated low expression and functionality of these factors in hepatocytes [8,13,18], concluding that this was the reason for the absence of an innate response to HBV infection. On the contrary, Verrier et al. [10] proposed that both proteins are expressed in hepatocytes and able to sense foreign DNA. Here we demonstrated that STING is expressed in hepatocytes including PHH, although to a reduced level compared to immune cells. cGAS RNA was detected in PHH, but was under the detection limit in HepG2-derived cell lines. Nevertheless, we show that PHH and HepG2 NTCP are able to sense DNA including HBV DNA, to some extend (Fig 5). Using THP-1 KO cell lines as a model to dissect the possible sensing pathway(s) involved (Fig 3), we showed that HT-DNA and HBV rcDNA sensing was totally abrogated by cGAS or STING KO, demonstrating that the cGAS/STING pathway is required for HBV rcDNA sensing and that other possible DNA sensors expressed in THP1 are not sufficient to sense these DNA species in the absence of cGAS or STING. However, we do not exclude that other DNA sensors such as IFI16, which was recently suggested to sense HBV cccDNA in the nucleus of hepatocytes [49], might act as cofactors of the cGAS/STING pathway. These data suggest that in hepatocytes, the cGAS/STING pathway is very likely responsible for the observed sensing of HBV rcDNA, even if cGAS and STING RNAs expression is low or below the detection limit of our assay. Furthermore, we demonstrated that the efficiency of the innate response to transfected DNA in hepatocytes is not optimal and can be improved by overexpressing cGAS and STING (Fig 5B). In summary, we propose a central role of the cGAS/STING pathway in HBV rcDNA sensing. We demonstrated that, in hepatocytes, this pathway is expressed at a relatively low level but is functional for foreign DNA sensing, although not fully efficient. Nevertheless, we revealed that PHH are able to respond to naked HBV rcDNA, suggesting that the low expression of the sensors cannot fully account for the absence of innate response to HBV infection. The possible inhibition of innate immune pathways by HBV as a way to escape the interferon response is still a matter of debate. The viral regulatory protein HBx has been described to interfere with MAVS [20, 26] leading to a reduced innate immune response to different stimuli in cells transfected with plasmids encoding HBx or HBV genome. However, the relevance of these observations in the context of HBV infection has not been investigated. Here we observed no innate immune response to HepG2-hNTCP or PHH infection with an HBV X-mutant (S1 Fig), neither ISG54 induction nor IRF3 phosphorylation, demonstrating that HBx is not responsible for the lack of the innate response to HBV in infected hepatocytes. Moreover, we showed that MAVS is not involved in sensing HBV nucleic acids, while cGAS and STING are (Fig 3).

Recent publications proposed that HBV does not inhibit the innate immune responses by suggesting that HBV infection does not alter the response to stimuli such as PolyIC, SeV, polydAdT [8,30,31], Rift Valley fever virus (RVFV) or MengoZn virus [31]. The majority of these stimuli trigger RNA-but not DNA-sensing pathways. In fact, only PolydAdT can activate DNA sensors but it can also activate the RIG-I pathway after transcription by RNA pol III. Therefore, these studies cannot rule out an inhibition of the DNA sensing pathway by HBV. However, Liu et al. suggested that HBV replication dampened STING-mediated innate signaling [29]. In addition, Verrier et al. [10] proposed that HBV infection decreases the expression levels of STING, cGAS and TBK1, but they did not reveal whether the functionality of the pathway was affected. On the contrary, Guo et al. [11] reported that induction of HBV replication in HepAD38 did not affect the response to foreign DNA but the authors did not study a genuine infection with HBV. So far, whether HBV inhibits the foreign DNA sensing pathway or not was still an open question. Here, we first demonstrated that IRF3 is not phosphorylated in HBV-infected hepatocytes (Fig 1C). This indicates that, if the absence of innate immune response to HBV is due to an active inhibition by the virus, this viral inhibition would target a step upstream of IRF3 phosphorylation. We then demonstrated that HBV infection does not alter IRF3 nuclear translocation in response to HT-DNA transfection (Fig 6). In addition, we observed no significant change in cGAS and STING expression levels in PHH infected with a high MOI of HBV (S6 Fig). Altogether our data point towards an absence of inhibition of the cGAS-STING DNA-sensing pathway by HBV.

In the absence of active inhibition, HBV evasion from the DNA-sensing pathway has to be passive. Since we observed that hepatocytes can sense and respond to naked HBV DNA, we hypothesize that shielding of viral rcDNA and replication intermediates into the viral capsids may be the reason why HBV DNA is not sensed upon infection. This hypothesis has been suggested in an immortalized mouse hepatocyte cell line model where HBV capsids are destabilized [50]. Nevertheless, the low efficiency of the cGAS/STING pathway in hepatocytes could be a second level of security for the virus, increasing the threshold for DNA recognition under conditions where small amounts of viral DNA are released in the cytoplasm from defective capsids.

In conclusion, we demonstrated that HBV does not induce an innate immune response in infected hepatocytes. We showed for the first time that HBV RNAs are not immunostimulatory. Furthermore, we indicated that different forms of HBV DNA can be sensed by the cGAS/STING pathway. This pathway is expressed at a low level but is active in hepatocytes and able to respond to naked HBV DNA. However, upon infection, the virus escapes the innate response without actively inhibiting the DNA-sensing pathway. The shielding of the viral DNA in the capsids is most likely the main mechanism allowing HBV to escape cGAS/STING sensing.

## Material and methods

*Details for cells, viruses, infections, lentiviral vectors and quantitative (reverse-transcription)-PCR are described in Supplementary methods*.

### Preparation and quantification of viral nucleic acids for transfection experiments

For monocyte-derived dendritic cells (MDDCs) and THP1 transfection experiments, total nucleic acids from HBV, Sendai Virus (SeV) and Modified vaccinia Ankara (MVA)-gfp viral stocks were extracted using the High Pure Viral Nucleic Acid Kit (Roche). When indicated, samples were digested with 1mg/ml of RNAseA for 1h at 37°C followed by 15 min inactivation at 72°C and addition of RNAseIN (Promega), or with DNase using the Turbo DNA-free kit (Ambion, Thermo Fisher Scientific) accordingly to manufacturer’s instructions.

Total RNAs from HepAD38 or HepG2 cells were extracted using the NucleoSpin® RNA Plus Kit (Macherey Nagel) and any DNA contaminants were digested with the Turbo DNA-free kit (Ambion, Thermo Fisher Scientific).

HBV replication intermediates were extracted from the cytoplasmic fraction of HepAD38 cells as described in [51] with minor modifications. Cells were incubated for 45 min on ice in hypotonic buffer (10 mM Hepes pH 8, 1,5mM MgCl2, 10mM KCl). 1% of NP-40 was then added and the samples were vortexed for 30s. Nuclei were pelleted by centrifugation at 6000 g at 4°C for 5 min. The supernatants (cytoplasmic fractions) were treated with 8U/ml of DNAseI in presence of 3,3mM of MgOAc for 90 min at 37°C in order to keep only encapsidated DNA. After proteinase K digestion, the DNA was extracted with phenol-chloroform and treated with RNAseA as described earlier.

HBV rcDNA, replication intermediates or MVA-gfp DNA copy numbers were determined by qPCR, HBV RNA and SeV RNA copy numbers were quantified using one-step reverse transcription qPCR (RT-qPCR) using specific primers and serial dilutions of standard DNA oligonucleotides (Supplementary Table 2). For RNA samples, the results are expressed as cDNA-equivalent since the copy number of the viral RNAs is assumed proportional to the respective specific cDNA quantification. RT-qPCRs and qPCRs were performed using QuantiTect SYBR Green RT-PCR Kit (Qiagen) respectively with and without RT on a LightCycler 480 (Roche).

### Transfections

Viral nucleic acids or HT-DNA (Sigma) were transfected into MDDCs or PMA-differentiated THP1 using Lipofectamine 2000 (Thermo Fisher Scientific) or in PHH and HepG2-hNTCP-derived cell lines using X-tremeGENE HP reagent (Merck) according to the manufacturer’s instructions.

For MDDC transfection experiments (Figs 2 and S2), the table 1 indicates the copy number/cell of viral DNAs (DNA copy/cell, determined by qPCR) or of viral RNAs (expressed as cDNA-equivalent copy/cell, determined by RT-qPCR). The indicated copy numbers/cell refer to the undiluted nucleic acids. SeV RNAs, MVA-gfp DNA and HBV DNA from viral particles were quantified from undigested nucleic acid preparations. HBV DNA replication intermediates were quantified from RNAse-digested samples. HBV RNA from viral particles or from total HepAD38 RNAs were quantified from DNase-digested samples to avoid any DNA contamination and a control without RT was performed.

### IRF3 nuclear translocation assays

HepG2-hNTCP overexpressing cGAS and STING (see Supplementary Methods for lentiviral transduction) were infected or not with HBV (MOI 10000) in presence of 2% DMSO. Four days post infection (dpi) the DMSO was removed. Six dpi, the cells were seeded on collagen-coated coverslides in 48-well plates and one day later transfected or not with 2 µg/well of HT-DNA. 16 hours post-transfection the samples were stained for IRF3 and HBV core (HBc) (see Supplementary Methods) and imaged with a Zeiss LSM 510-META Confocal laser scanning microscope.

## Acknowledgements

The authors thank Nina Hein-Fuchs and Esther Schnellbächer for experimental support.

## Supplementary methods

### Cells and ethics statements

HepG2-hNTCP cells are derived from HepG2 cells and have been stably transfected with HBV receptor, the sodium-taurocholate cotransporting polypeptide (NTCP) [1]. HepG2.2.15 cells are derived from HepG2 cells and have been stably transfected with HBV genome [2]. HepG2 H1.3deltaX cells are derived from HepG2 cells and contain the stable integration of a 1.3-fold HBV genome carrying premature stop codon mutations in both 5’ and 3’ parts of the HBx open reading frame [3]. HepG2-hNTCP, HepAD38, HepG2 H1.3deltaX were maintained in Dulbecco’s modified eagle medium (DMEM) containing 10% fetal calf serum (FCS) and 2 mM L-Glutamine (complete DMEM). HepaRG-hNTCP are derived from HepaRG and overexpress NTCP. They were grown in William’s E medium supplemented with 10% FCS, 2 mM L-hNTCP, HepAD38, He^-5^ M hydrocortisone hemisuccinate, and 5 μg/ml insulin.

THP-1 cells deficient for cGAS (CGAS), STING (TMEM173) and MAVS (MAVS) were previously described [4]. THP-1 cells were cultivated in RPMI supplemented with 10% fetal calf serum (FCS) and 2 mM L-Glutamine. For differentiation, 40 ng/ml of phorbol 12-myristate 13-acetate (PMA) were added for 48 hours.

Cell lines were authentified as described in the Supplementary Table 1 and regularly tested to exclude any mycoplasma contamination using a PCR based method with the PCR Mycoplasma Test Kit II from AppliChem.

**Supplementary Table 1.** Cell lines, suppliers and authentification methods.

| Name | Citation | Supplier | Authentication test method |
| --- | --- | --- | --- |
| HepG2 | Fig. 4 | Gift from Prof | Cell line authentication performed by |
|  |  | S.Urban, Heidelberg | eurofins genomics, using Applied Biosystems™ AmpFLSTR™ Identifiler™ Plus PCR Amplification Kit with 16 markers |
| HepG2-hNTCP | Fig. 1, 4, 5, 6, S1, S3, S4 | Gift from Prof S.Urban, Heidelberg (Ni, Gastroenterology 2014) | Cell line authentication performed by eurofins genomics, using Applied Biosystems™ AmpFLSTR™ Identifiler™ Plus PCR Amplification Kit with 16 markers |
| HepaRG-hNTCP | Fig. 4 | Gift from Prof S.Urban, Heidelberg | Multiplexion cell contamination assay (Heidelberg, Germany) as described [1] |
| HepaD38 | Fig. 2, S2 (for HBV RNAs and replication intermediate extraction) | Gift from Prof E Hild, Langen | Release of HBV particles (qPCR + infectivity test) |
| THP1 | Fig. 3, 4 | Gift from Prof V. Hornung, Munich | Cell line authentication by eurofins genomics on 24.03.2017 |
| THP1 ΔSTING | Fig. 4 | Gift from Prof V. Hornung, Munich | Western blot |
| THP1 ΔcGAS | Fig. 4 | Gift from Prof V. Hornung, Munich | Western blot |
| THP1 ΔMAVS | Fig. 4 | Gift from Prof V. Hornung, Munich | Western blot |

Primary human hepatocytes (PHH) were isolated from liver specimens resected from patients undergoing partial hepatectomy. Approval from the local and national ethics committees (Ethic committee from the Goethe University of Frankfurt, agreement number 343/13, and the Hannover medical school, agreement number 252-2008 and informed consent from patients were obtained. PHH were isolated with a two-step perfusion method and cultured as described previously [5]. Alternatively, cryopreserved plateable PHH were purchased from Thermo Fisher Scientific. PHH were maintained in Williams’ Medium E supplemented with Serum-free Hepatocyte Maintenance Supplement Pack (Thermo Fisher Scientific CM4000) according to the provider’s instruction.

Cryopreserved human Kupffer cells were purchased from Thermo Fisher Scientific and handled according to the provider’s instructions.

Monocyte-derived dendritic cells (MDDCs) or monocyte-derived macrophages (MDMs) were generated from human peripheral blood mononuclear cells (PBMC) using respectively 25 U/ml GM-CSF (Leukine®Sargramostim, Genzyme) and 800 U/ml IL-4 (PeproTech) or 50 U/mL GM-SCF. The PBMCs were isolated from human buffy coats of anonymous blood donors purchased from the German Red Cross Blood Donor Service Baden-Württemberg Hessen.

### HepG2-hNTCP overexpressing cGAS and STING

HepG2-hNTCP were transduced with pLX304-based lentiviral vectors encoding V5-tagged cGAS and selected with 20 µg/ml of blasticidin. For each experiment, the selected cells were freshly transduced with pLX304-based lentiviral vectors encoding V5-tagged STING and used 6 days post transduction for experiments.

### Hepatitis B Virus production and infection

Preparation of Hepatitis B Virus wild type (HBVwt) inoculum from HepAD38 cells or HBV deficient for HBx (HBV X-) from HepG2 H1.3deltaX was performed as previously described (Rivière et al 2015). Viral particles were tittered by qPCR quantification of viral RC-DNA with the primers HBV_RC_F and HBV_RC_R (Supplementary Table 2). For infection, HepG2-hNTCP and PHH were seeded on collagen-coated plates. HBV infections were performed in presence of 4% Polyethylene glycol 8000 and 2% DMSO unless otherwise indicated. When indicated, HBV entry inhibitor MyrcludexB [6] (1 µM) was added. 24 hours after viral inoculation, the medium was changed and complete DMEM supplemented with 2% DMSO was added.

### Other viruses and viral vectors

Sendai virus, strain Cantell, was provided by G. Kochs, Medical Center-University of Freiburg, Germany.

The DNA virus MVA-gfp based on Modified vaccinia Ankara was kindly provided by Gerd Sutter [7, 8].

### Quantitative reverse transcription PCR (RT-qPCR)

For RT-qPCR analysis, total RNAs were extracted from cell pellets using NucleoSpin® RNA Plus Kit (Macherey Nagel). RT-qPCRs were performed on a LightCycler 480 (Roche) using QuantiTect SYBR Green RT-PCR Kit (Qiagen) and specific primers (Supplementary Table 2). For analysis of HBV RNAs, primer sets that specifically amplify the 3.5 kb, 2.4 kb and 2.1 kb HBV RNAs (HBV RNA 3 F, HBV RNA 3 R, Supplementary Table 2) were used. Unless otherwise stated, Ribosomal Protein L13 A (RPL13A) was used as a reference gene to normalize the samples. For each sample, the relative amount of RNA was calculated using the formula 2-deltaCt with delta Ct = (Ct gene of interest – Ct RPL13A). When indicated, relative change to the control condition was calculated using the formula 2-delta delta Ct with delta delta Ct = delta Ct sample – delta Ct control.

For Figure 4, the expression levels of the genes of interest and of 2 reference genes, RPL13A and TATBP, were calculated using a standard curve generated with serial dilutions of THP1 RNA. For each sample, the values of the genes of interest were normalized to the geometric mean of the 2 reference genes. The limit of detection for each sample and each gene of interest was calculated as the RNA amount measured in the last detectable standard dilution of THP1 for the gene of interest divided by the geometric mean of the expression levels of the 2 reference genes for each sample.

**Supplementary Table 2.**
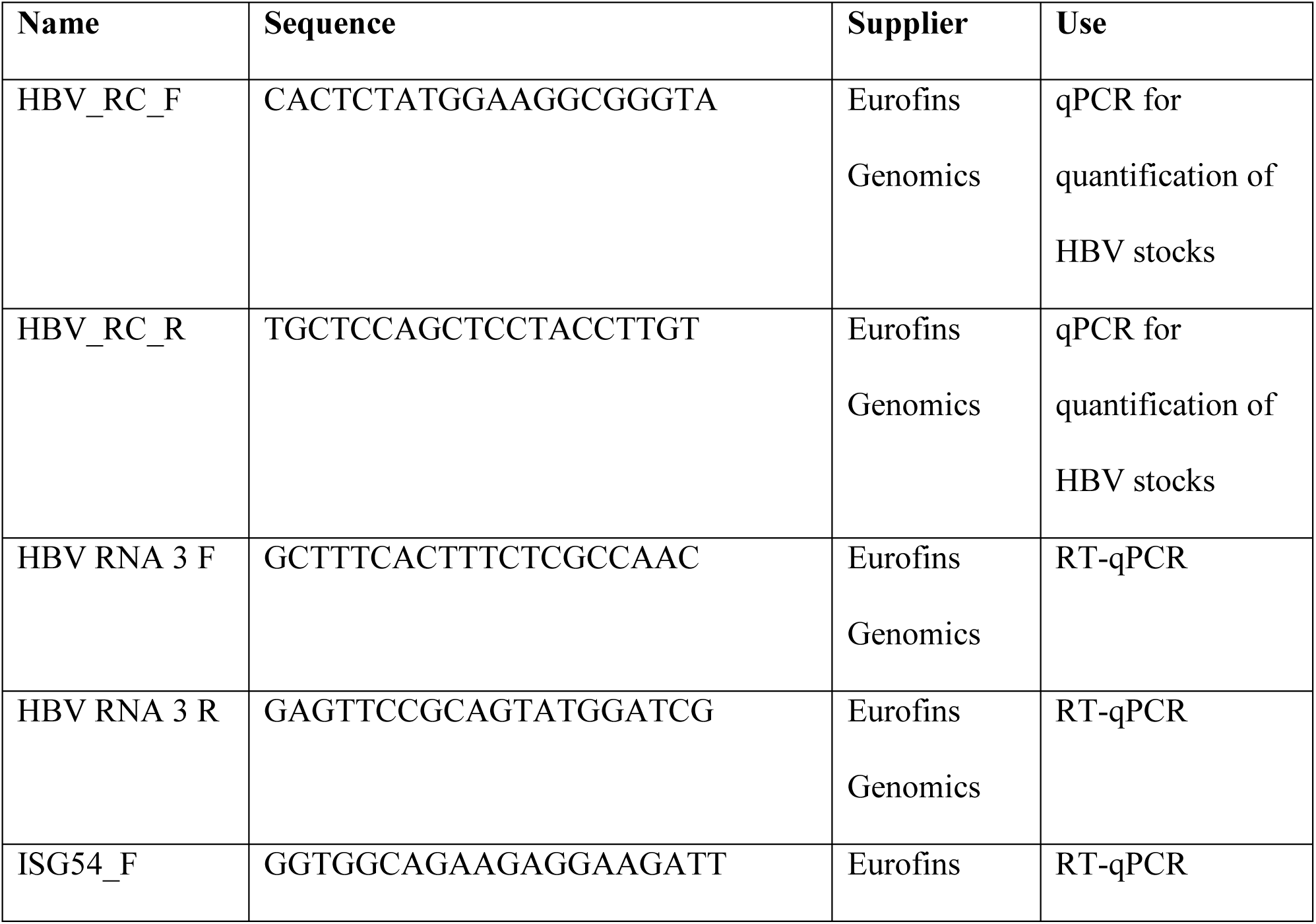
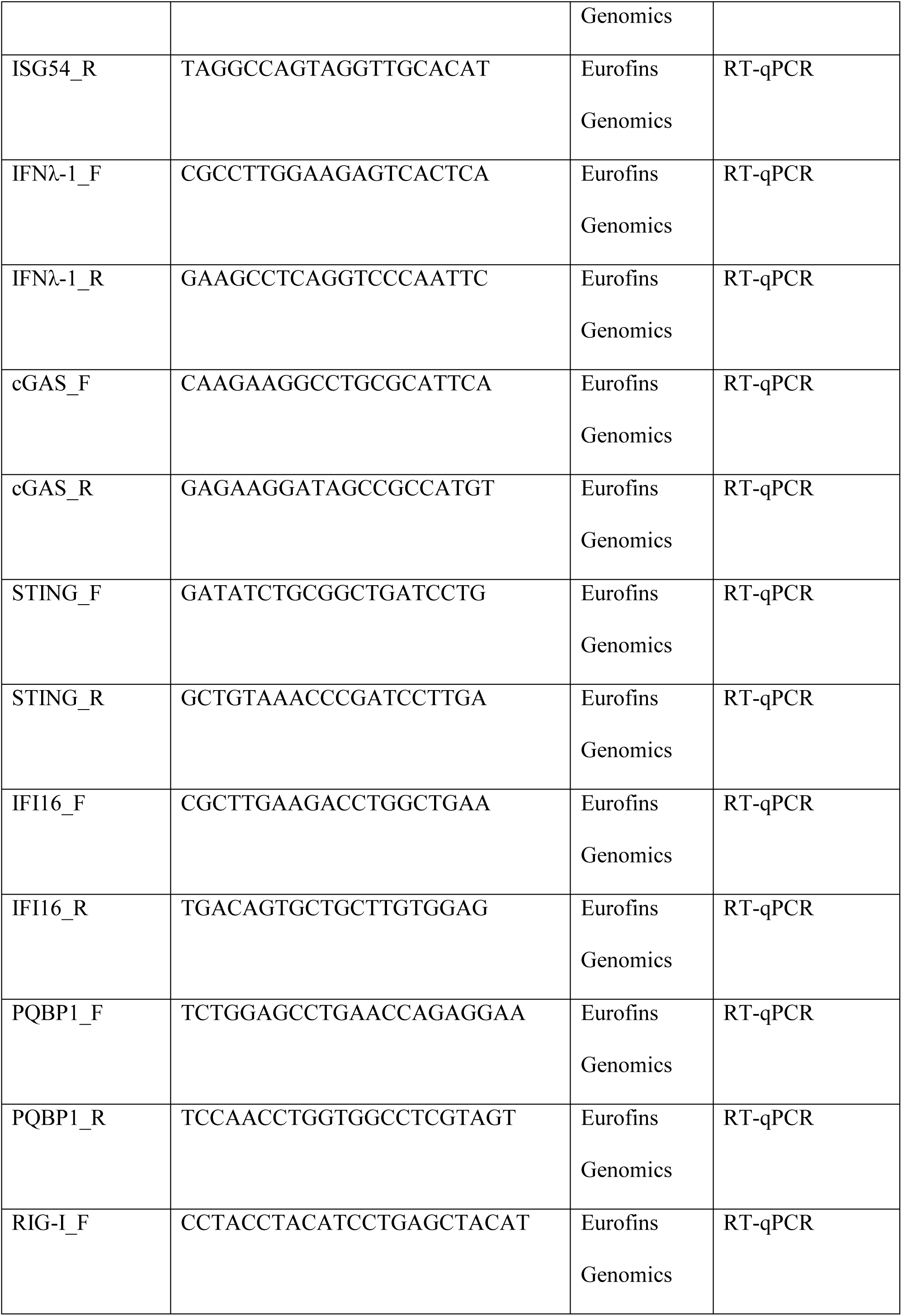
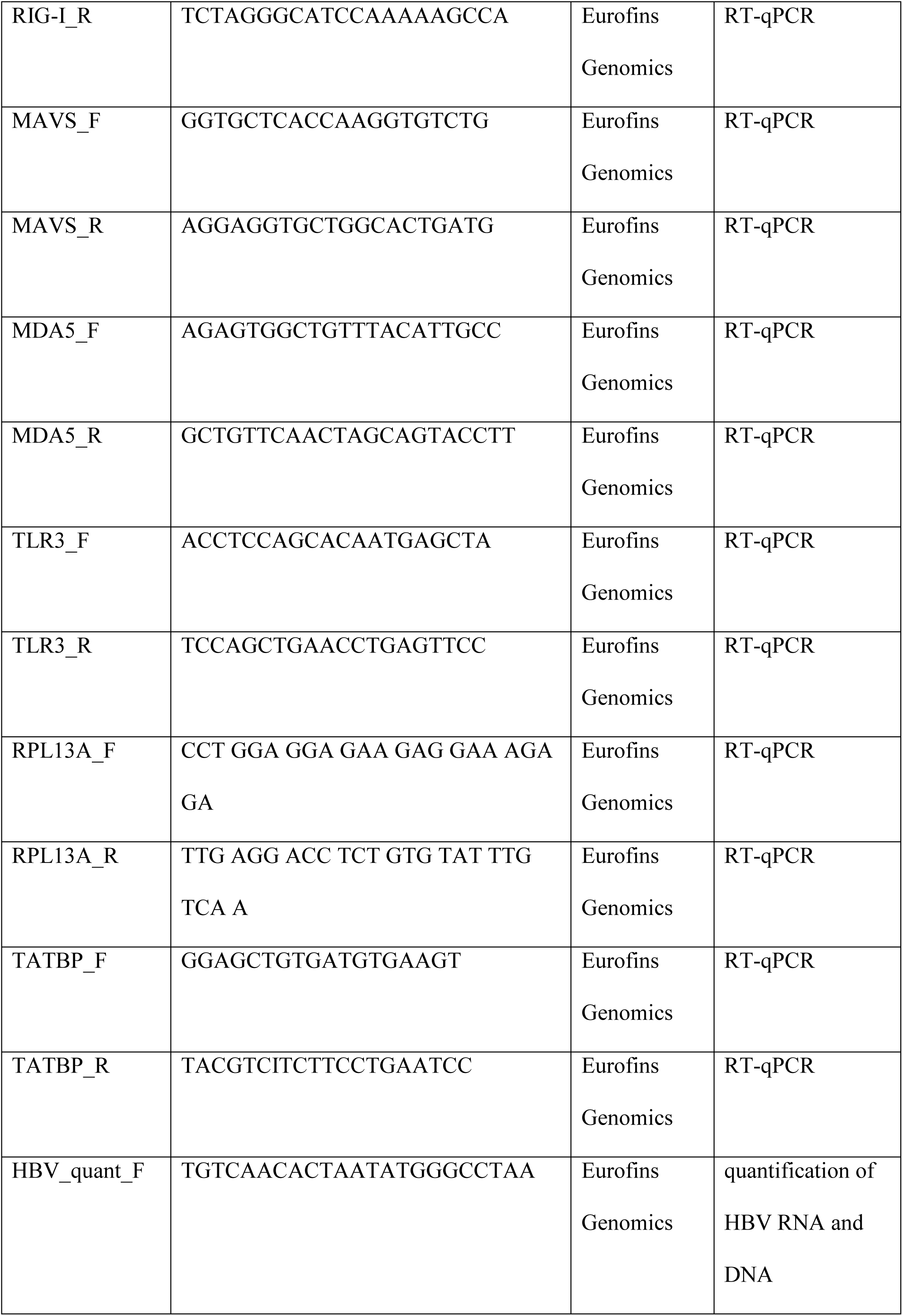
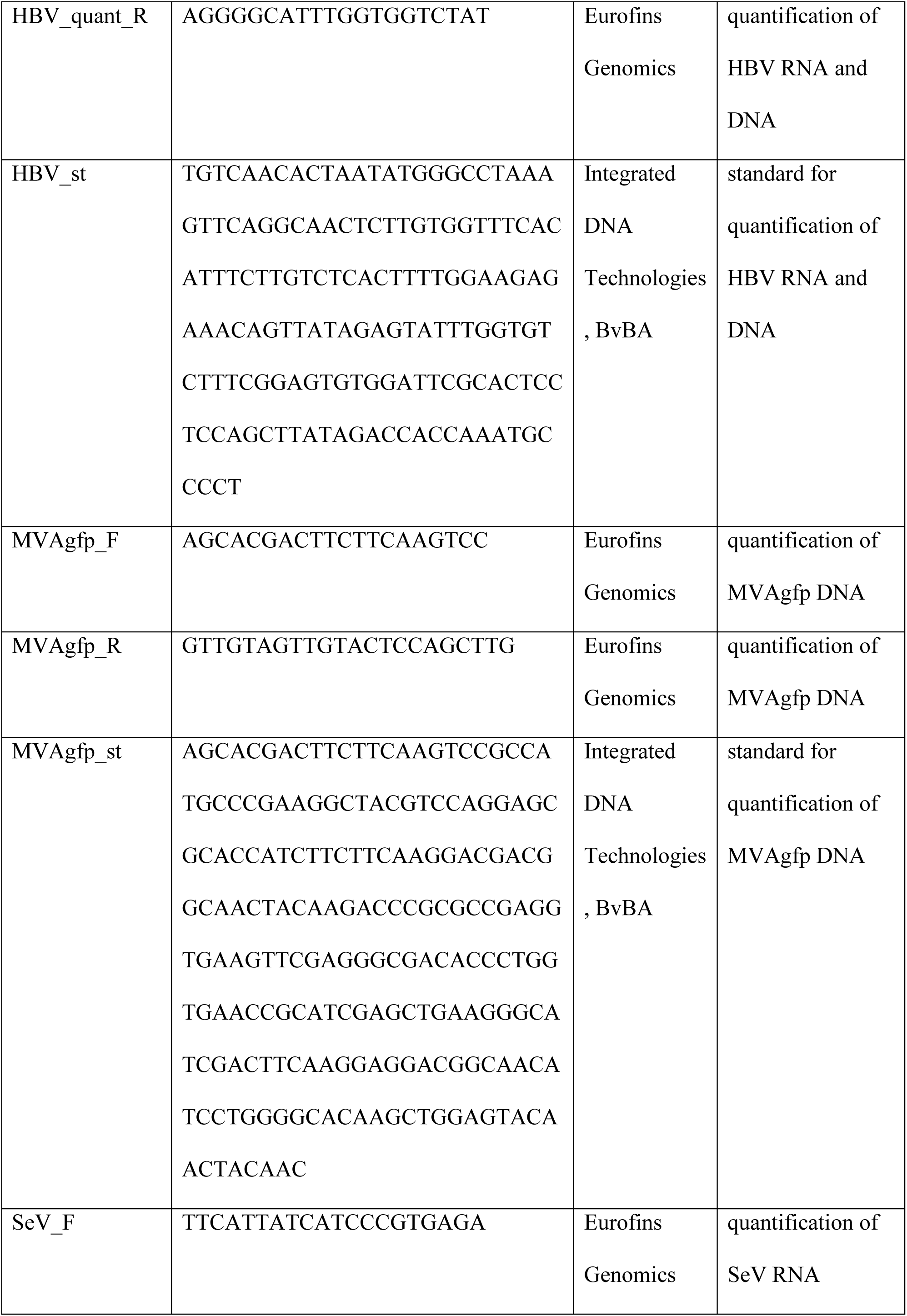
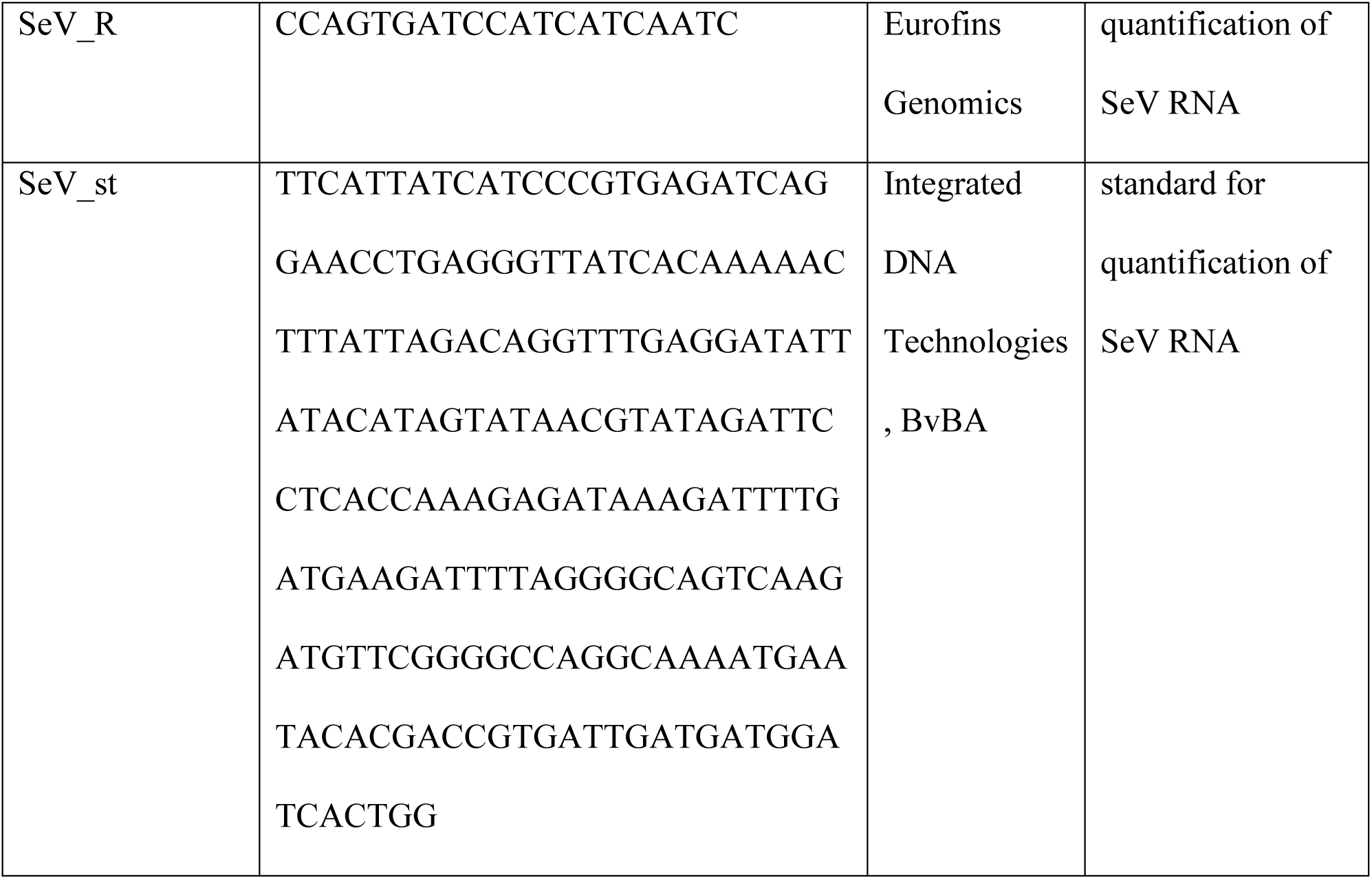
Oligonucleotides.

### Reagents

2’3’ cGAMP was purchased from Invivogen and used at a final concentration of 2µg/ml in cell supernatant. HT-DNA was purchased from Sigma-Aldrich.

### Western blots

Cell samples were lysed using a TritonX-100 based Lysis Buffer (100 mM NaCl, 10 mM EDTA, 20 mM TRIS pH7.5, 1% TritonX-100, 1% Sodiumdesoxycholate). Proteins were separated by SDS-PAGE and transferred to a Nitrocellulose membrane. Specific proteins were detected using the antibodies listed in the Supplementary Table 3. Secondary antibodies conjugated to the horseradish peroxidase (GE Healthcare) were used for chemiluminescence detection.

**Supplementary Table 3.**
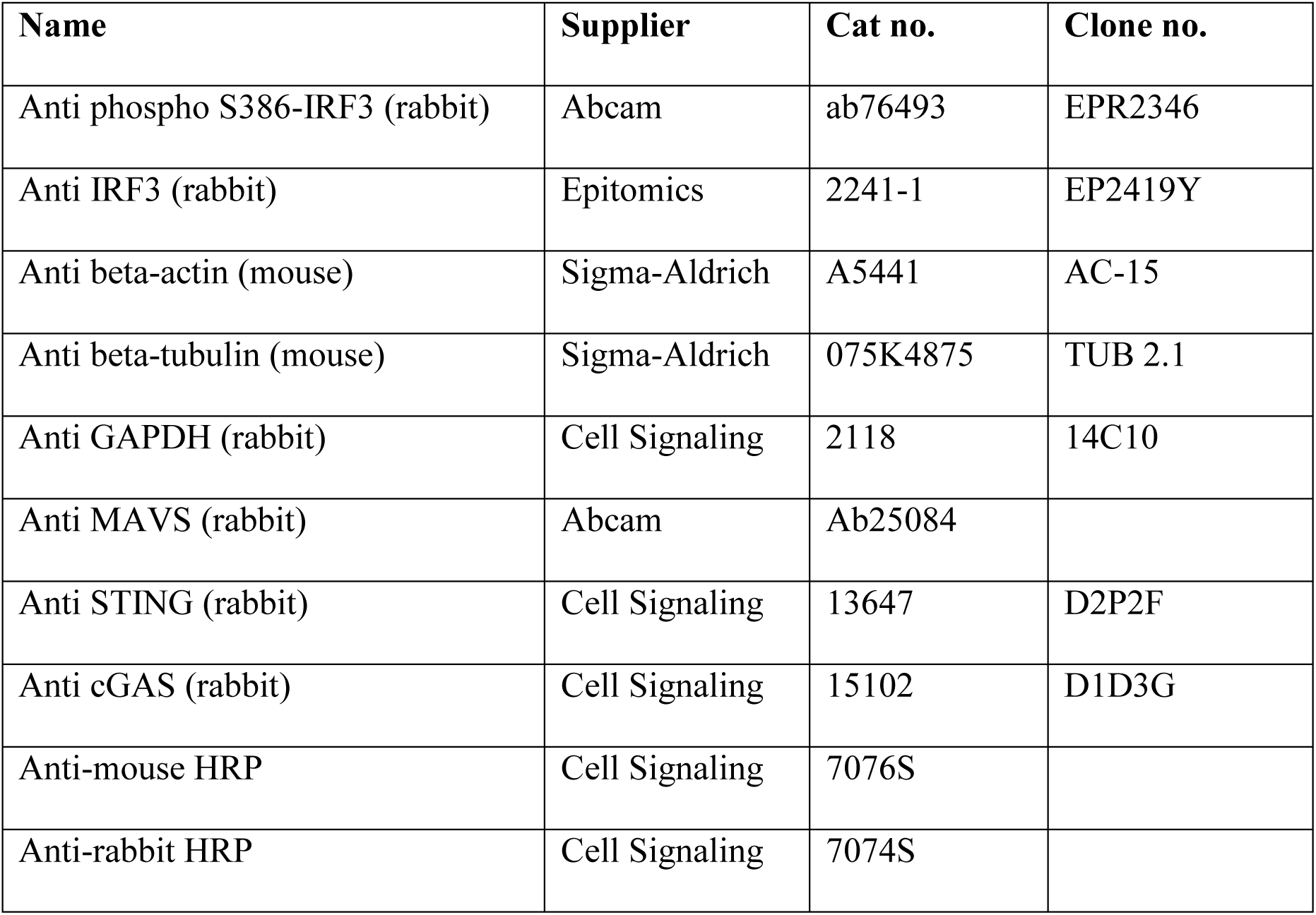
Antibodies used for Western blots.

### Immunofluorescence

Cells were fixed with 4% paraformaldehyde for 15 min, washed 3 times with PBS, permeabilized with 0,25% Triton-X100 for 7 min at room temperature and blocked for 1 hour in PBS-0,1% Tween (PBST) containing 5% of bovine serum albumine (BSA). The coverslides were then incubated for 2 hours at room temperature with the primary antibodies, washed 3 times in PBST, and incubated for 1 hour with the secondary antibodies (Supplementary Table 4). The coverslides were washed 3 times in PBST and the DNA was stained with Hoechst reagent.

**Supplementary Table 4.** Antibodies used for immunofluorescence.

| **Name** | **Citation** | **Supplier** | **Cat no.** | **Clone no.** |
| --- | --- | --- | --- | --- |
| Anti IRF3 (rabbit) | IRF3<br>translocation<br>assay, Fig.6 | Cell Signaling | 11904S | D6I4C |
| anti HBc (mouse) | IRF3<br>translocation<br>assay, Fig.6 | gift from Prof. S.<br>Urban, Heidelberg |  | M312 |
| anti-rabbit Alexa<br>Fluor 488 | IRF3<br>translocation<br>assay, Fig.6 | Invitrogen | A11008 |  |
| anti-mouse Alexa<br>Fluor 555 | IRF3<br>translocation<br>assay, Fig.6 | Thermo Fischer<br>Scientific | A21422 |  |
| Anti-HBc (rabbit) | Fig. S6 | Dako | B0586 |  |
| Goat anti-rabbit<br>IgG(H+L), Alexa<br>Fluor 546 | Fig. S6 | ThermoFisher | A-11010 |  |

### Microscope image acquisition

For the IRF3 translocation assay (Fig.6) the samples were mounted with Mowiol medium. They were imaged using a Zeiss LSM 510-META Axiovert 200M Confocal laser scanning microscope, with a 40x objective (EC “Plan-Neofluar” 40x/1,30 Oil DIC) and an Axiocam camera with the acquisition software ZEN 2009, Version: 5.5.0.451.

Imaging of the MVA-gfp-infected HepG2-hNTCP (Supplementary Fig. 3) was performed in the culture plates with cell medium using a Nikon Eclipse TS100 inverted microscope with a 10x objective (Achromat, 10x/0,25 Ph1 DL) and a Nikon DS-Vi1 camera, with the acquisition software NIS-Elements F 4.30.00 (Build 1020) 64 bit.

For HBcAg staining in PHH (Supplementary Fig. 6), stained cells were imaged in PBS without mounting medium using a LEICA DM IRB microscope with a 20x objective (N PLAN L 20*/0.40 CORR PH 1) and a HAMAMATSU camera with the acquisition software HoKawo 2.12.

All microscope acquisitions were performed at room temperature. Contrast/brightness of the images were adjusted with Fiji Image J (1.52p).

## Supplementary figures

**S1 Fig:**
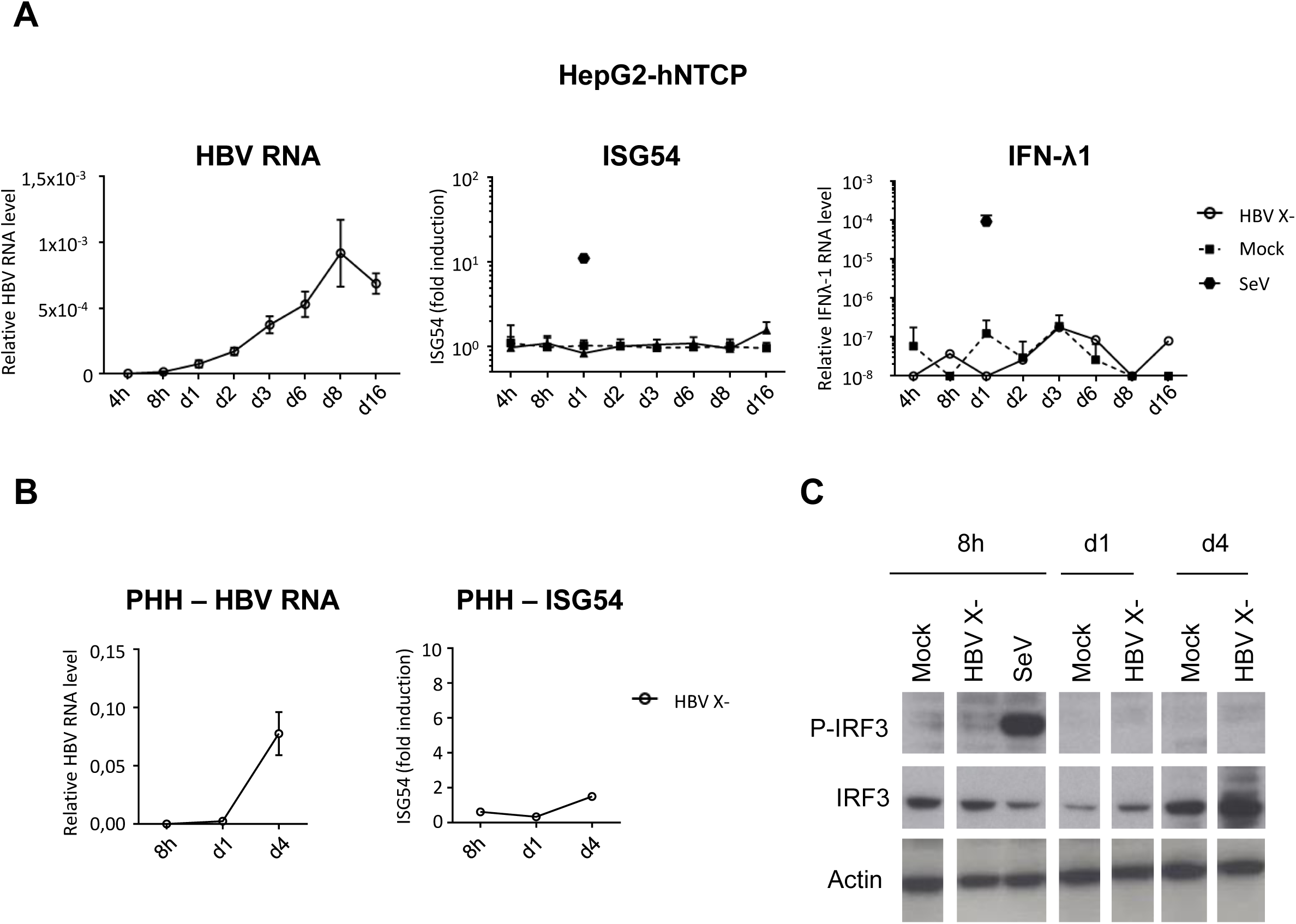
No innate immune response to HBV infection in the absence of HBx. HepG2-hNTCP (A) or PHH (B, C) were infected with HBV X- (MOI 100) or with SeV (MOI 0,13). HBV, ISG54 or IFN lambda-1 RNAs were measured by RT-qPCR (A, B). Relative expression levels to the reference gene RPL13A were calculated. For ISG54, fold induction to the respective mock-control is shown for each time point. Average and SEM of 3 independent experiments in triplicates (HepG2-hNTCP) or 1 representative experiment in triplicate (PHH) are shown. (C) Phospho- or total IRF3 protein levels were analyzed by Western blot in infected PHH.

**S2 Fig:**
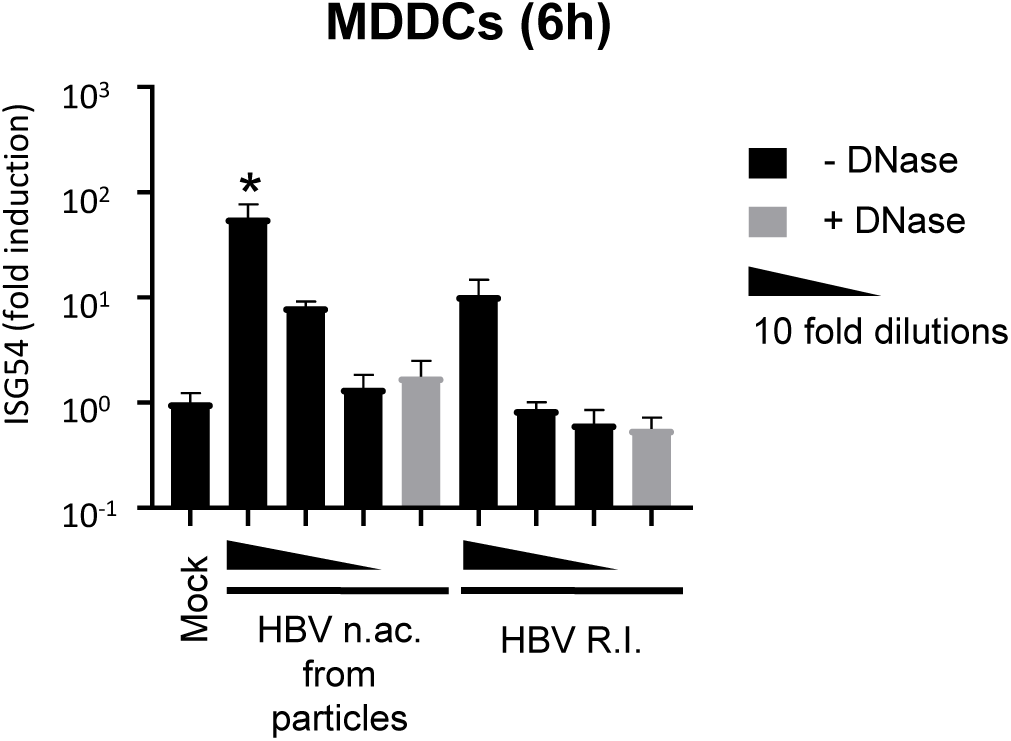
HBV rcDNA as well as replication intermediates are immunostimulatory. MDDCs were transfected with 5000 copies/cell and 10-fold serial dilutions of total nucleic acids (n.ac.) from HBV particles or of HBV DNA replication intermediates (HBV R.I.) from HepAD38 cytoplasm. When indicated, the nucleic acids were DNAse-digested (light grey) prior transfection (using the same volume as for 5000 copies/cell of undigested nucleic acids). ISG54 mRNA was quantified by RT-qPCR 6h post-transfection. Average and SEM of 3 technical replicates of at least 2 donors are shown. ****p<0,0001, ***p<0,001, **p<0,01 and *p<0,05 ‘(Kruskal and Wallis test with Dunn’s test).

**S3 Fig:**
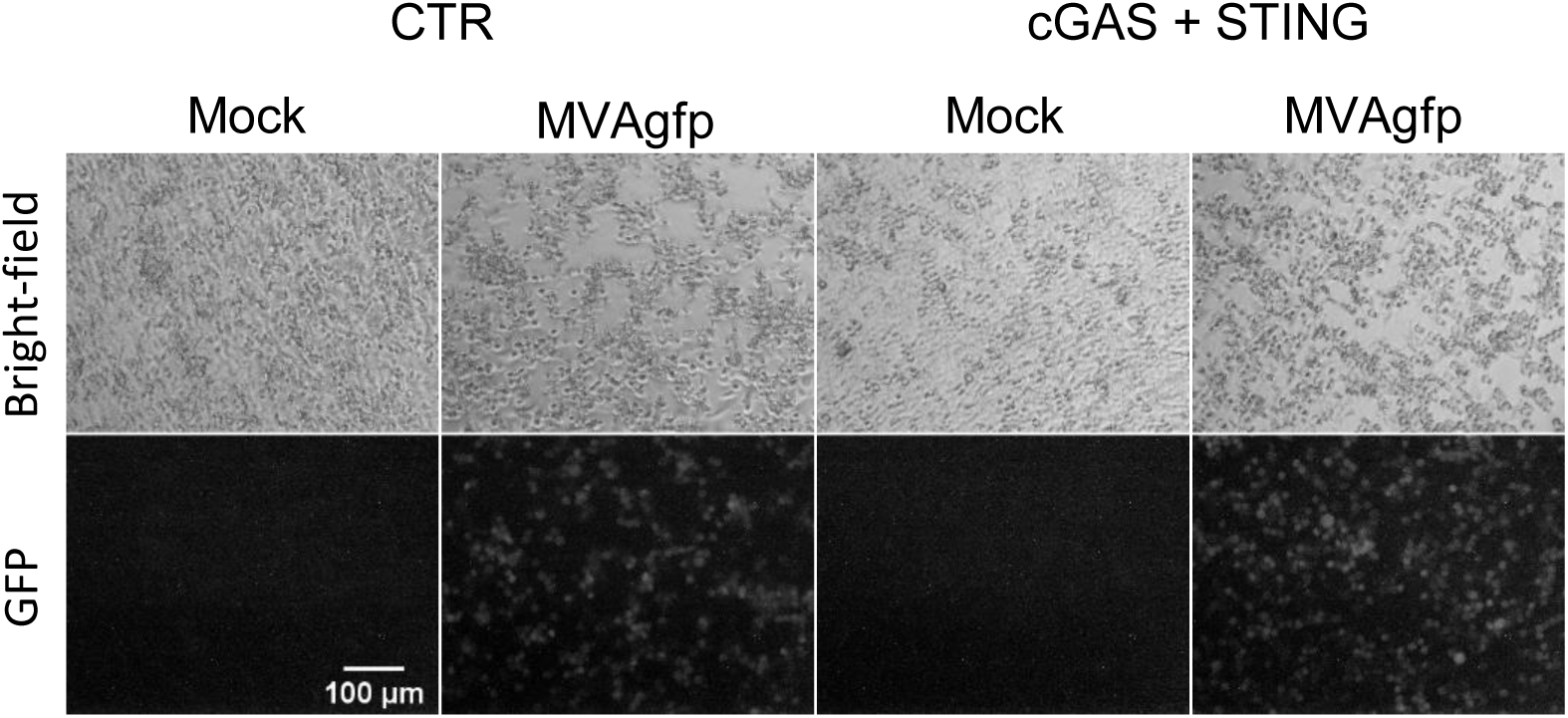
Infection control for MVAgfp infection of HepG2-hNTCP. Gfp expression in the MVAgfp-infected HepG2-hNTCP CTR or cGAS+STING cells used in Fig.5, 20h post infection.

**S4 Fig:**
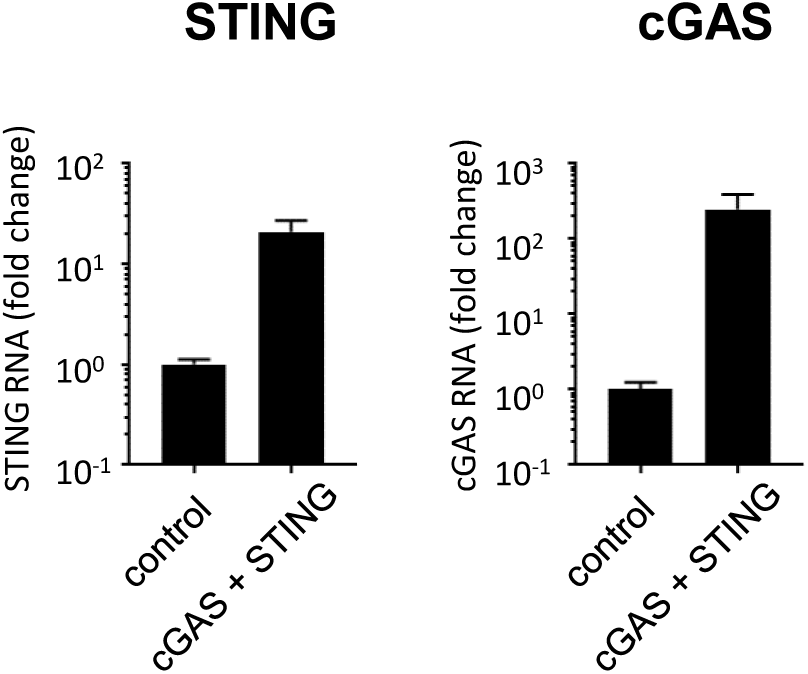
cGAS and STING overexpression in HepG2-hNTCP. The expression level of cGAS and STING mRNA in HepG2-hNTCP transduced with a control lentiviral vector or with lentiviral vectors expressing cGAS and STING used in Fig 5B was determined by RT-qPCR. The graph shows the average and SEM of 2 experiments performed in technical triplicates.

**S5 Fig:**
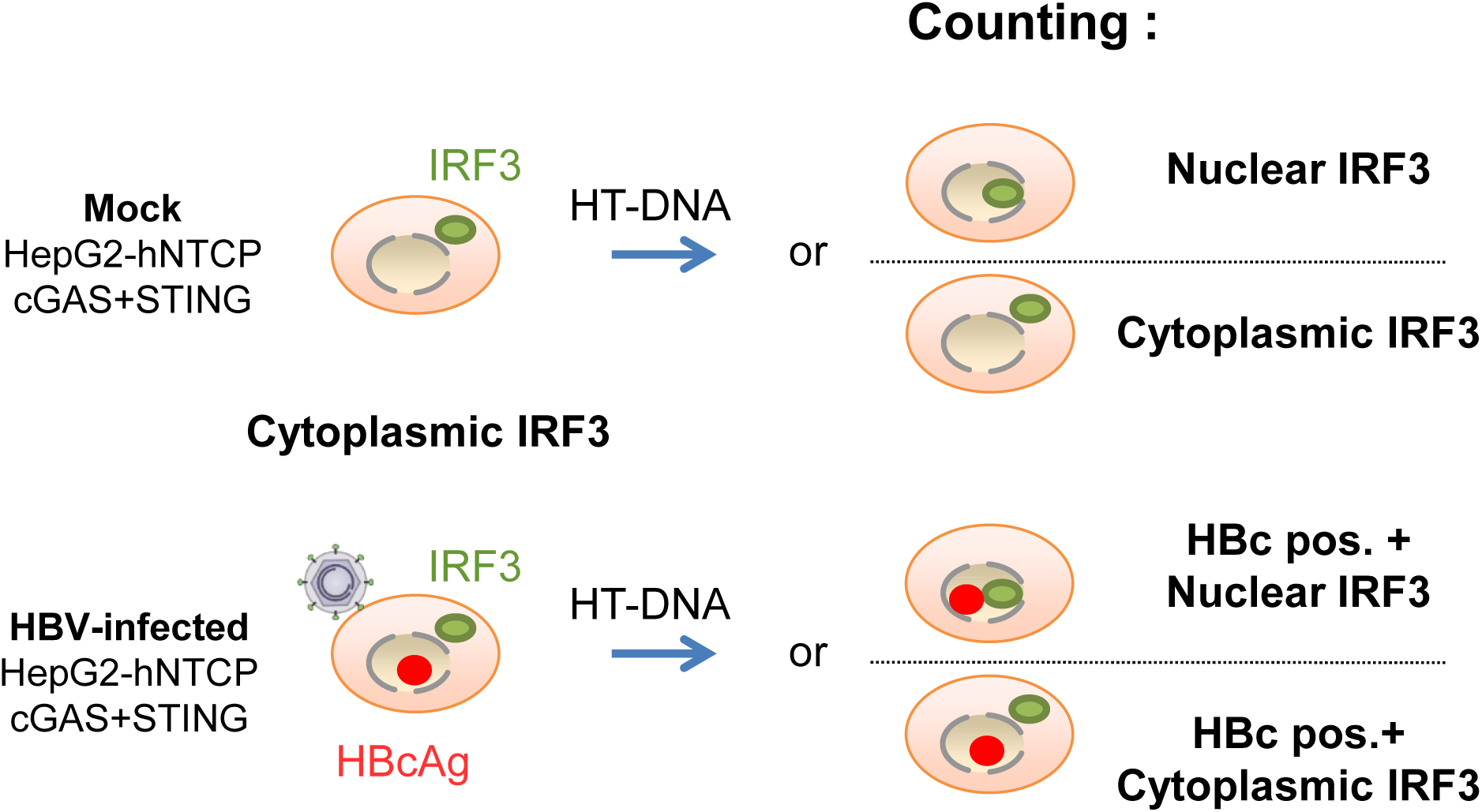
Experimental design for Fig 6. HepG2-hNTCP overexpressing cGAS and STING were infected or not with HBVwt (MOI 10000). 7 dpi, the cells were transfected with HT-DNA to stimulate the DNA sensing pathway. 16 hours post transfection the cells were fixed and immunostained for IRF3 (Alexa 488, green), HBc (Alexa 555, red) and DNA (Hoechst, blue). The percentage of cells harboring a nuclear staining for IRF3 in the mock-infected cells after HT-DNA transfection or in the HBc positive cells from the HBV-infected and HT-DNA-transfected samples was counted and is indicated in the graph Fig. 6B.

**S6 Fig:**
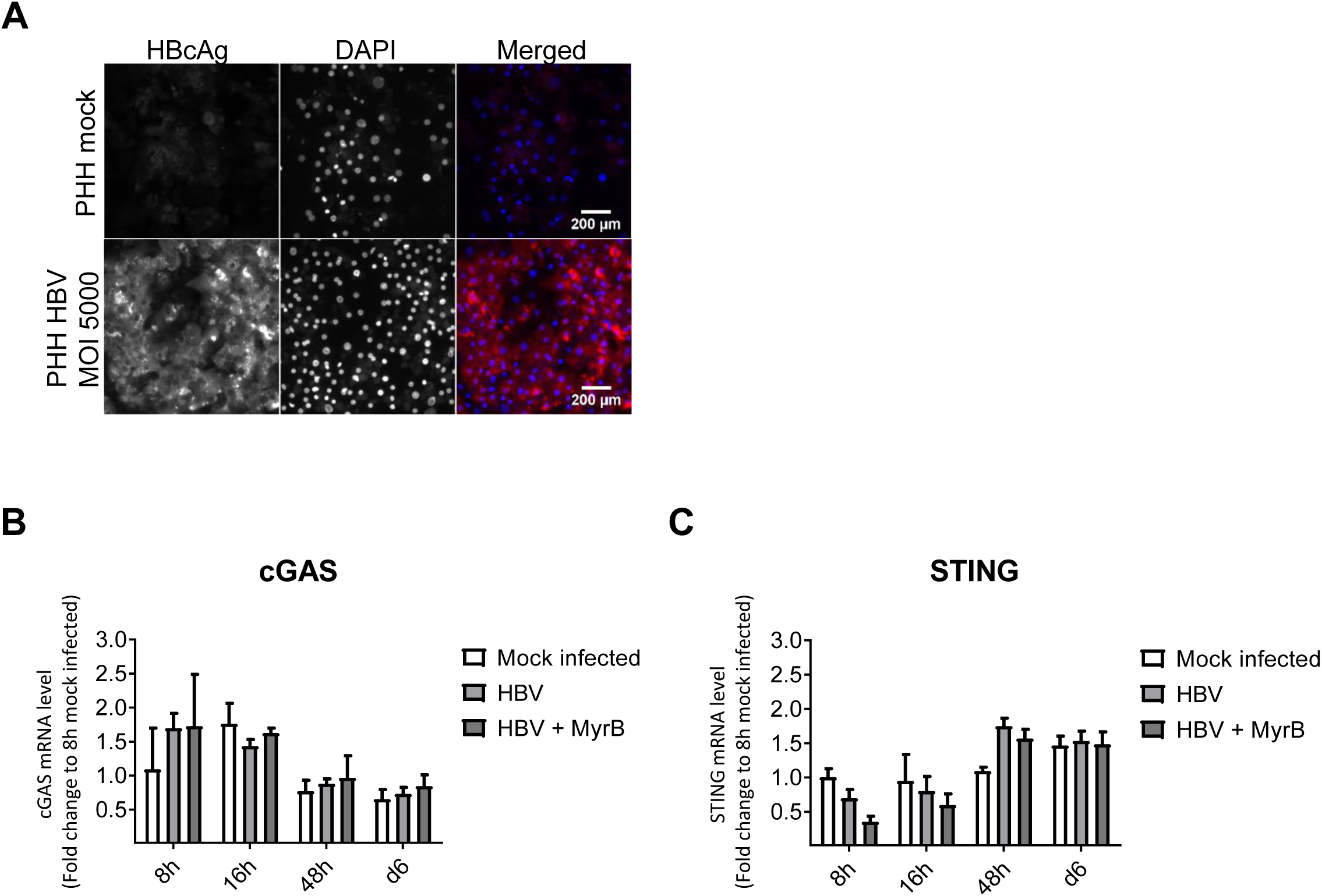
HBV does not affect the expression level of cGAS or STING. PHH were infected with HBVwt (MOI 5000). (A) Immunofluorescence staining of HBcAg (red) in mock- or HBV-infected PHH 7 days post infection. cGAS (B) or STING (C) mRNA levels were quantified by RT-qPCR at the indicated times post infection. The HBV entry inhibitor Myrcludex B (MyrB; 1µM) was added as a control. The graph represents the average and standard error of the mean (SEM) of 3 technical replicates of at least 2 donors.

**S7 Fig:**
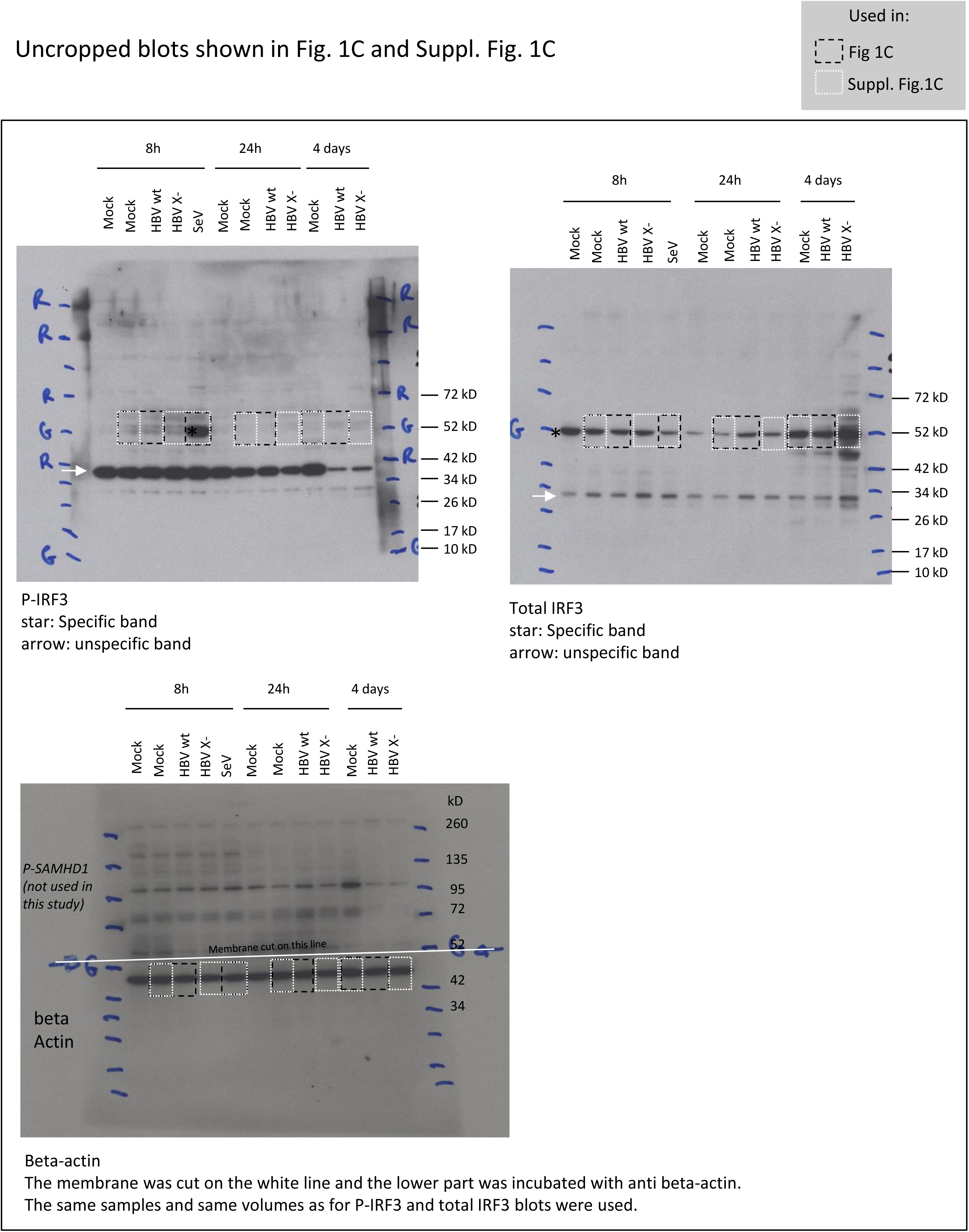
Uncropped western blots. The uncropped western blots shown in Fig 1C, and S1C are presented in this figure.

**S8 Fig:**
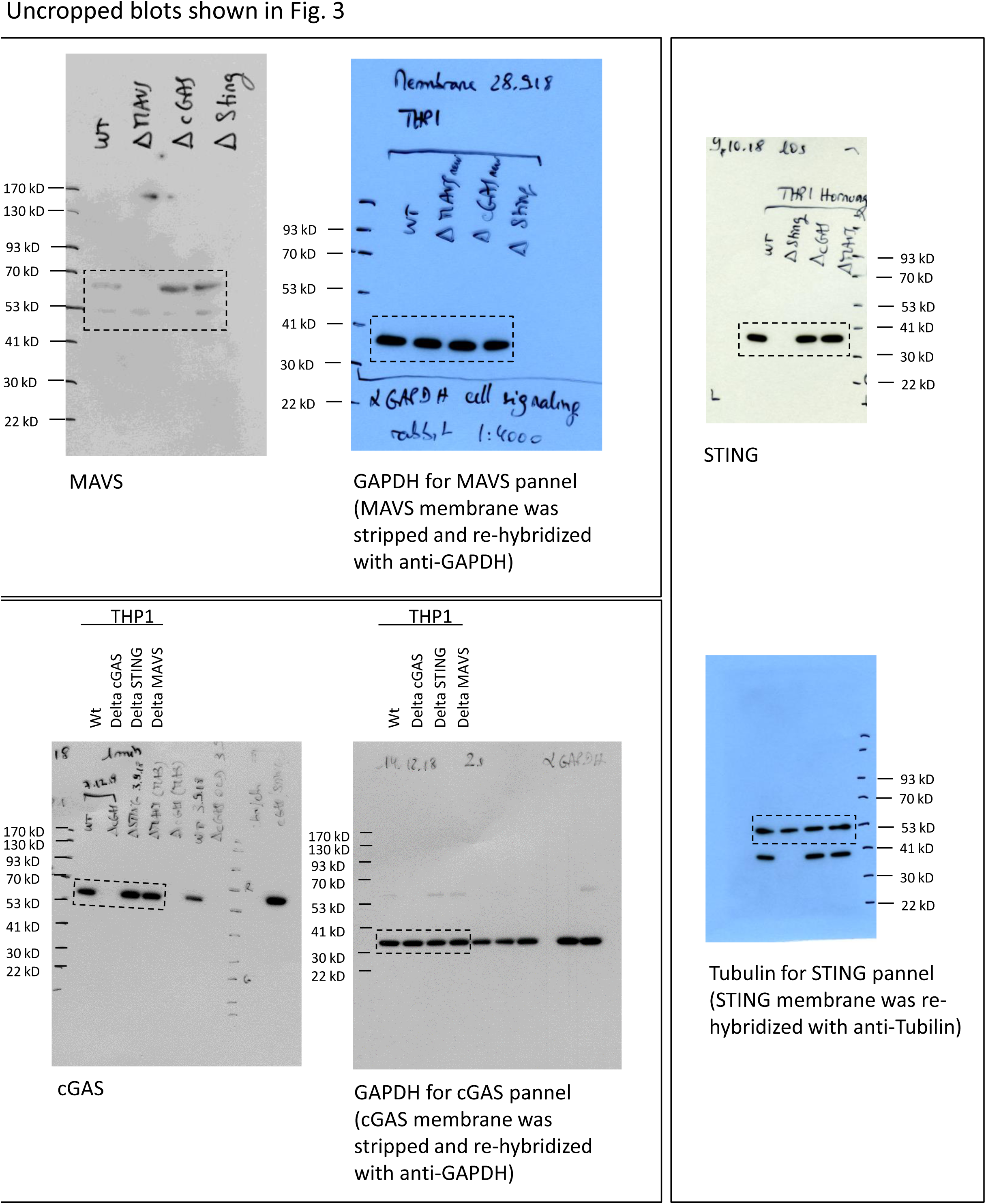
Uncropped western blots. The uncropped western blots shown in Fig 3 are presented in this figure.

